# Abnormal expression of splicing regulators RBFOX and NOVA is associated with aberrant splicing patterns at the Neurexin-3 gene in a monogenic autism spectrum disorder

**DOI:** 10.1101/2025.10.02.680092

**Authors:** Pedro K. K. Forti, Lilian L. Depieri, Lucas A. Giorgiani, Breno B. Hernandes, João G. Bueno, Gustavo D. Verçosa, Israel C. Vasconcelos, Mario H. Bengtson, Marcelo F. Carazzolle, Antonio P. Camargo, Fabio Papes

## Abstract

Autism spectrum disorders are diseases characterized by a combination of cognitive, behavioral and neurological symptoms. A complex interplay between environmental factors and a multitude of genetic determinants, most of them composed of low-risk variants, impose difficulties in understanding the molecular and cellular underpinnings of these conditions. In some cases, autistic patients have been shown to display alterations in splicing patterns of several genes, but the extent to which this phenomenon is common in the context of these multifactorial disorders is unknown, nor is it known if monogenic cases of autism also display dysregulation of splicing. Moreover, very few studies have investigated the causal links between splicing alterations in specific genes and the phenotypic characteristics of the neural tissue in autism patients. In this study, we have focused on a monogenic type of autism caused by haploinsufficiency of the Transcription Factor 4 gene, known as Pitt-Hopkins Syndrome. We show that neurons and organoids derived from patients with this disease have altered expression of splicing regulators in the NOVA and RBFOX families, accompanied by aberrant splicing patterns in a substantial number of genes involved in neural tissue development and synapse organization. We focused on a gene encoding a member of the Neurexin Family, the splicing of which normally leads to the production of transcript variants coding for both transmembrane and secreted protein isoforms. In Pitt-Hopkins Syndrome neurons, we detected an aberrant splicing pattern that results in lower expression of the secreted isoform and its corresponding transcript, a phenomenon that may explain why the neural tissue in these patients have decreased electrical activity through impaired synapse organization. Our data shed light on the role splicing regulation plays in a monogenic type of autism.

## INTRODUCTION

RNA splicing plays a critical role in regulating transcription and producing the wide diversity of transcripts and protein isoforms that characterize the eukaryotic cell. Alternative splicing results in the joining of exons and exclusion of introns in different configurations, being classified as exon skipping (SE), intron retention (RI), alternative 5’ or 3’ splice site (A5SS or A3SS), and mutually exclusive exon (MXE) splicing events^1^. RNA splicing is tightly regulated by RNA-binding proteins, several of which are important to neurogenesis, axon guidance, synaptogenesis, and synaptic physiology during neurodevelopment^2^.

Alternative splicing is very prevalent in the human brain, particularly during neuronal differentiation^3^, affecting events such as synaptic function and stability^4^. Aberrant splicing patterns at genes involved in synaptic plasticity such as *NLGN3*, *NLGN4*, *NRXN1*, *SHANK3*, and *CADPS2* are present in individuals with autism-spectrum disorders^5^, which has been taken as a suggestion that several of these genes may have a putative role in the molecular mechanisms driving autism.

Out of the many RNA-binding proteins regulating splicing, two families have been linked to autism in several studies – RNA Binding Fox-1 Homolog (RBFOX) and Neuro-Oncological Ventral Antigen (NOVA) proteins. RBFOX family members RBFOX1, RBFOX2, and RBFOX3 are expressed in neurons^6^, regulating alternative splicing patterns particularly during early neurogenesis and for synaptic plasticity (RBFOX1 and RBFOX2) or in mature neurons (RBFOX3)^7^. RBFOX1 controls the skipping of neuron-specific exons in many genes, particularly in those related to synapse formation, like *NRCAM* and *GRIN1*, which are dysregulated in samples of autistic children^8^. Likewise, NOVA1 has been shown to bind introns flanking neuron-specific exons, leading to exon inclusion or skipping^9^. In the brains of autistic children or individuals with other types of neurodevelopmental disorders, some studies have identified downregulation of RBFOX and NOVA family members^1,10^, and several direct RBFOX and NOVA target genes are involved in synaptic transmission or ion channel regulation^1,11^. Additionally, RBFOX variants have been associated with several conditions characterized by neuropsychiatric symptoms^12^.

Despite recent advances in understanding the role played by the dysregulation of RNA splicing regulators in the pathogenesis of autism, little is known about how mutations in specific genes causing autism-spectrum disorders lead to RBFOX and NOVA alterations and aberrant splicing patterns in downstream genes. This gap of knowledge prompted us to investigate a monogenic type of autism. Here, we have studied the occurrence of aberrant alternative splicing patterns in a monogenic autism disease called Pitt-Hopkins Syndrome (PTHS; MIM 610954)^13–16^. PTHS is caused by de novo heterozygous mutations in the Transcription Factor 4 gene (*TCF4*; OMIM 602272), which encodes a bHLH transcription factor expressed in the brain^17–20^ and important for neurodevelopment, neural lineage commitment, and neuronal physiology^21–26^. PTHS individuals exhibit severe cognitive deficit, motor delay, hypotonia, respiratory abnormalities, autistic behaviors, chronic constipation, and distinctive facial features^27,28^. Interestingly, variants of the *TCF4* gene have also been associated with schizophrenia, bipolar disorder, post-traumatic stress disorder, and major depressive disorder^29–33^, suggesting that part of the underlying molecular mechanisms may be common between these conditions and autism. Our choice to study aberrant splicing in PTHS is due to its monogenetic nature, which allows us to dissect its pathological molecular mechanisms and more precisely correlate them with the cellular abnormalities found in the PTHS neural tissue.

We have previously generated and analyzed neural progenitor cells, neurons, and brain organoids from patients with PTHS to analyze the molecular phenotypes of the diseased neural tissue^34,35^. We showed that PTHS neural progenitors cultured using traditional 2D techniques and neural progenitors in PTHS organoids display a deficit in proliferation and severely diminished differentiation rate into neurons. Strikingly, we detected that PTHS organoids representative of the cerebral cortex are aberrant in size and structure, containing a higher percentage of neural progenitor cells and fewer cortical neurons of many subtypes, including deep-layer and superficial-layer excitatory neurons; moreover, PTHS organoids representative of the subpallial region exhibit a deficient content of cortical-type GABAergic interneurons^34^.

Importantly, PTHS neurons in 2D culture and the neural network of PTHS organoids are characterized by severely impaired electrical activity, as judged by patch-clamp and multi-electrode array assays, respectively. Additionally, assessment of neuronal activity via FOS expression revealed that FOS+ neurons are fewer in PTHS organoids, concomitant with a lower expression of excitatory neuron’s synaptic markers vGLUT1 and vGLUT2^34^, suggesting that synapse formation is altered in this type of autism.

In this paper, we studied the expression of RBFOX and NOVA splicing regulators in PTHS neurons, showing that the lower expression of those regulators is accompanied by distinct splicing profiles in a range of genes, in comparison with neurons of normal genotype. These aberrant splicing patterns affect several genes coding for proteins important for synapse physiology, including Neurexin genes. We found that the Neurexin-3 gene (*NRNX3*) displays an aberrant pattern of splicing involving an exon skipping event, where PTHS neurons have relatively lower abundance of a transcript variant that includes an exon leading to a premature stop codon and producing a soluble secreted form of Neurexin. On the other hand, neurons from donors of normal genotype exhibit lower abundance of the other transcript variant (excluding the exon), which produces a transmembrane Neurexin thought to be involved in synapse formation and stability^36^. Our findings show the novel information that a severe form of autism is accompanied by altered splicing of a gene encoding a synaptic component, leading to an imbalance between different Neurexin neuronal isoforms, which may contribute to the PTHS neural tissue’s impaired electrical activity.

## RESULTS

### Altered expression of splicing regulators *RBFOX1/2/3* and *NOVA1* in Pitt-Hopkins Syndrome neurons in 2D culture

We have previously shown that the neural tissue in Pitt-Hopkins Syndrome (PTHS) is characterized by poor proliferation of neural progenitor cells and impaired differentiation of cortical neurons, in both traditional 2D neuronal cultures and in patient-derived cortical organoids^34^. We also detected lower neural activity in PTHS neurons, which could be in part due to impaired synaptogenesis. Importantly, we analyzed the gene expression profiles of both PTHS neurons and organoids via bulk RNA sequencing followed by differential gene expression analysis^34^ to identify dysregulated genes and pathways that could potentially be causally implicated with these phenotypes. These transcriptomic data were obtained from 4 PTHS patients harboring heterozygous mutations in the *TCF4* gene, causally related to the disease^34^.

Interestingly, we noticed that several entries in the list of differentially expressed genes between PTHS samples and same-sex parental controls code for RNA splicing regulators, such as RNA Binding Fox-1 Homolog 1 (RBFOX1), RBFOX2, and RBFOX3, as well as Neuro-Oncological Ventral Antigen 1 (NOVA1) and NOVA2 (Supplementary Table S1)^34^. Even though *NOVA2* is also downregulated in PTHS neurons in 2D culture, its overall expression levels across parental controls and patients are lower than *NOVA1* (Supplementary Table S1). Therefore, we decided to focus our efforts on understanding the role played by *NOVA1* and *RBFOX1/2/3* in triggering aberrant splicing patterns in PTHS.

In fact, several patient-parent pairs exhibited lower expression of *RBFOX1/2/3* and *NOVA1* genes in PTHS neurons. For *RBFOX1*, the fold-changes (in log base 2) of expression in the comparison between patient-control pairs #1, #2, and #3 are −4.95, −2.03, and −5.1, respectively (Fig. 1a-d); for patient #4, the expression is low, comparable to the levels displayed by the other 3 patients, although a matching parental control was unavailable in this case (Supplementary Table S1). For *RBFOX2*, the fold-changes between patient-control pairs #1, #2, and #3 are - 2,26, −1.79, and −1,61, respectively (Fig. 1a-d and Supplementary Table S1). *RBFOX3,* the most studied splicing regulator in neurons, coding for the neuronal marker NeuN, exhibits fold-changes of −10.9, −2,93, and −5,14, for pairs #1, #2, and #3, respectively (Fig. 1a-d and Supplementary Table S1). Finally, *NOVA1* is also significantly less expressed in PTHS samples as compared to parental controls for patient-control pairs #1, #2, and #3 (Fig. 1a-d), and its expression is low in patient #4 (Supplementary Table S1).

**Figure 1:**
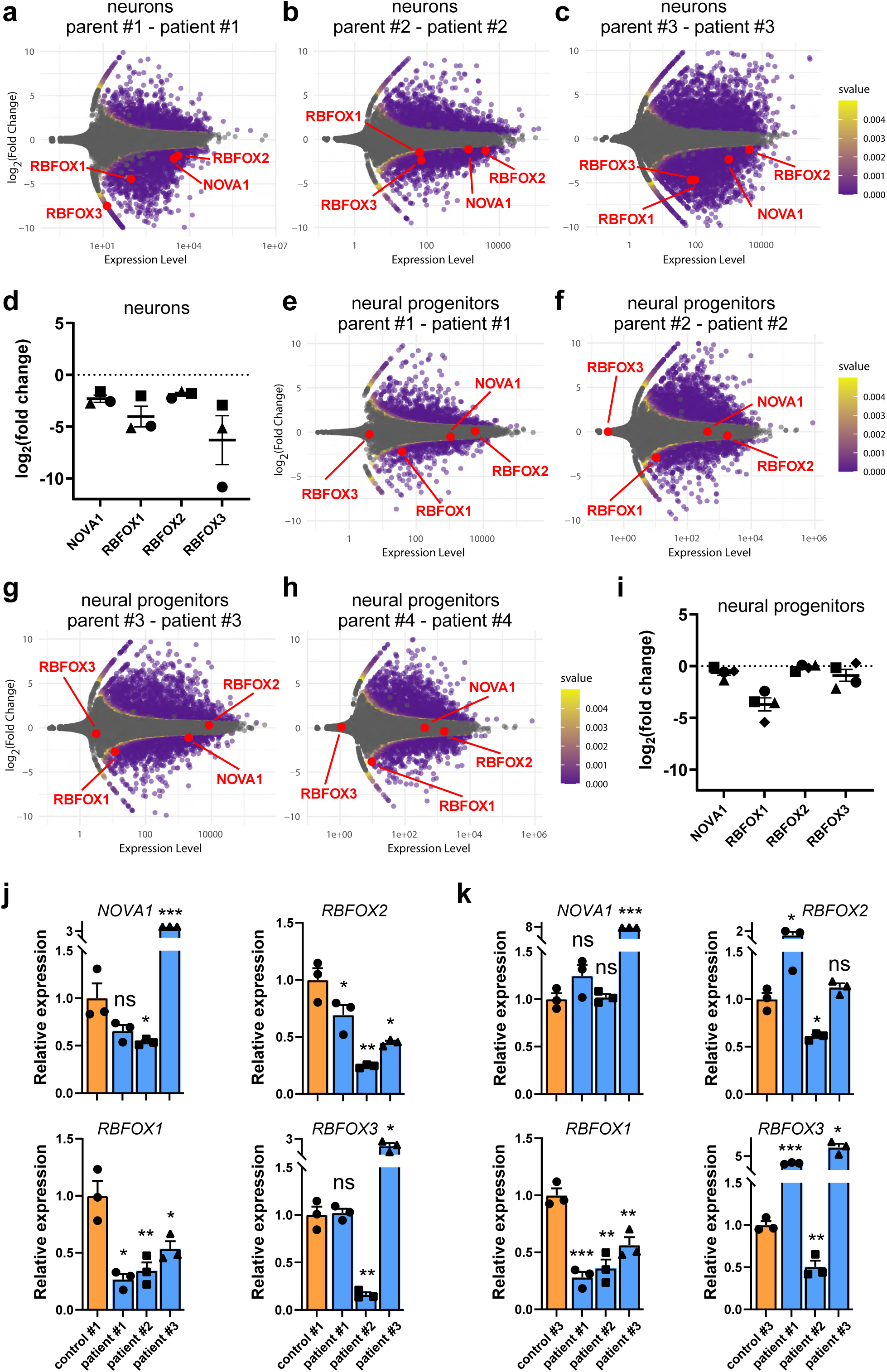
Lower expression of splicing regulators NOVA1 and RBFOX1/2/3 in neurons derived from Pitt-Hopkins Syndrome (PTHS) patients. (a-c) MA plots showing differential gene expression in neurons in 2D culture in the comparison between different pairs of control and PTHS subjects – pair #1 in (a), pair #2 in (b) and pair #3 in (c). See Supplementary Table S6 for patient information. Each parent-patient comparison is shown separately. Gray dots are genes that are not statistically significantly differentially expressed between control and PTHS. Colored dots are statistically significant DE genes, and the color indicates adjusted *p*-value (*s*-value). Red dots represent genes coding for splicing regulators NOVA1, RBFOX1, RBFOX2, and RBFOX3. The *x* axes represent average normalized expression across all libraries. Fold-changes (in log scale, base 2) in the *y* axes indicate the contrast between expression levels in the PTHS patient and the corresponding control (negative values indicate lower expression in PTHS neurons). (d) Summary of fold changes (in log scale, base 2) of *NOVA1* and *RBFOX1/2/3* gene expression for each patient-parent pair. Symbols indicate different pairs, as shown in Supplementary Table S6. (e-h) MA plots showing differential gene expression in neural progenitor cells in 2D culture in the comparison between different pairs of control and PTHS subjects – pair #1 in (e), pair #2 in (f), pair #3 in (g), and pair #4 in (h). (i) Summary of fold changes of *NOVA1* and *RBFOX1/2/3* gene expression in neural progenitor cells. Notice the lack of change in expression of some of these splicing regulators in neural progenitors. (j,k) Relative expression levels for *NOVA1*, *RBFOX1*, *RBFOX2*, and *RBFOX3* genes, as determined from RT-qPCR experiments, in iPSC-derived neuronal cultures. Three patients per graph, with mean expression normalized to 1 in control #1 (j) or control #2 (k). N = 2 technical replicates per sample. Bar graphs represent mean + SEM. ns, not statistically significantly different, \**p*<0.05, \*\**p*<0.01, \*\*\**p*<0.001; two-sample Welch’s t-test assuming unequal variances.

The impaired expression of *RBFOX1/2/3* and *NOVA1* in PTHS neurons seems to be specific to this neural cell category, as PTHS neural progenitor cells do not show a diminishment in the expression levels of such genes: only the expression of *RBFOX1* is lower in patient-parent pairs #1 to #4, while the other genes show average fold-changes close to zero (Fig. 1e-i). Importantly, the overall expression abundance of these genes is somewhat lower in neural progenitor cells as compared to neurons, as judged by the metric of expression abundance TPM (transcripts per million reads) (Supplementary Table S1).

Next, we performed independent neuronal differentiation experiments to quantify the expression of *NOVA1* and *RBFOX1/2/3* in PTHS neurons via reverse transcriptase-coupled quantitative PCR (RT-qPCR) in comparison with two controls of normal genotype. Similar to the transcriptomic data obtained from bulk RNA-Seq experiments in 2D cultured neurons, *RBFOX1* expression levels were −1.9, −1.5, and −0.9-fold lower in patients #1, #2, and #3 in the comparison with control #1, or −1.8, −1.5, and −0.8-fold lower in patients #1, #2, and #3 in the comparison with control #3 (Fig. 1j). Similarly, *RBFOX2* expression levels were also lower in patients #1, #2, and #3 in the comparison with control #1 and in patient #2 in the comparison with control #3 (Fig. 1j). On the other hand, *RBFOX3* expression was only found to be lower in patient #2, and NOVA1 expression was lower in patients #1 and #2 in the comparison with patient #1 (Fig. 1j).

In combination, the RNA-Seq and qPCR results indicate that splicing regulators in the NOVA and RBFOX families have lower expression levels in neurons of certain PTHS individuals cultured using traditional 2D techniques. It is noteworthy that *RBFOX1* is consistently downregulated in PTHS neurons across all subjects in our cohort.

### Complex dysregulation in the expression of *RBFOX1/2/3* and *NOVA1* in Pitt-Hopkins Syndrome cortical and subpallial organoids

Because there was a difference in expression abundances for *RBFOX1/2/3* and *NOVA1* between two different neural cell types in our bulk RNA-Seq data, we sought to explore the expression of these genes in single-cell transcriptomic data from PTHS patient-derived cortical and subpallial organoids, which are complex 3D structures able to model the events that happen during early neurodevelopment in vivo and. These models are more representative of the molecular characteristics of neuronal cells than traditional 2D cultures^34,37,38^. Because the organoids contain a more diverse set of neurons and the interactions between different cell types may influence the dysregulation of *NOVA* and *RBFOX* expression, we reasoned that the organoids would more faithfully recapitulate the changes in expression occurring in vivo. In organoid single-cell RNA sequencing (scRNA-Seq) libraries previously described by our group^34^, we could annotate six neural cell subpopulations, including progenitor cells, intermediate progenitors, and neurons^34^ in two distinct differentiation lineages that produce either glutamatergic or GABAergic neurons (Fig. 2a). In cortical organoids (CtOs), the predominant differentiation lineage leads to glutamatergic neurons (Fig. 2b), while subpallial organoids (sPOs) generate mostly GABAergic cells (Fig. 2b), matching the differentiation profile found in the cortex and subpallial brain regions in vivo^39,40^.

**Figure 2:**
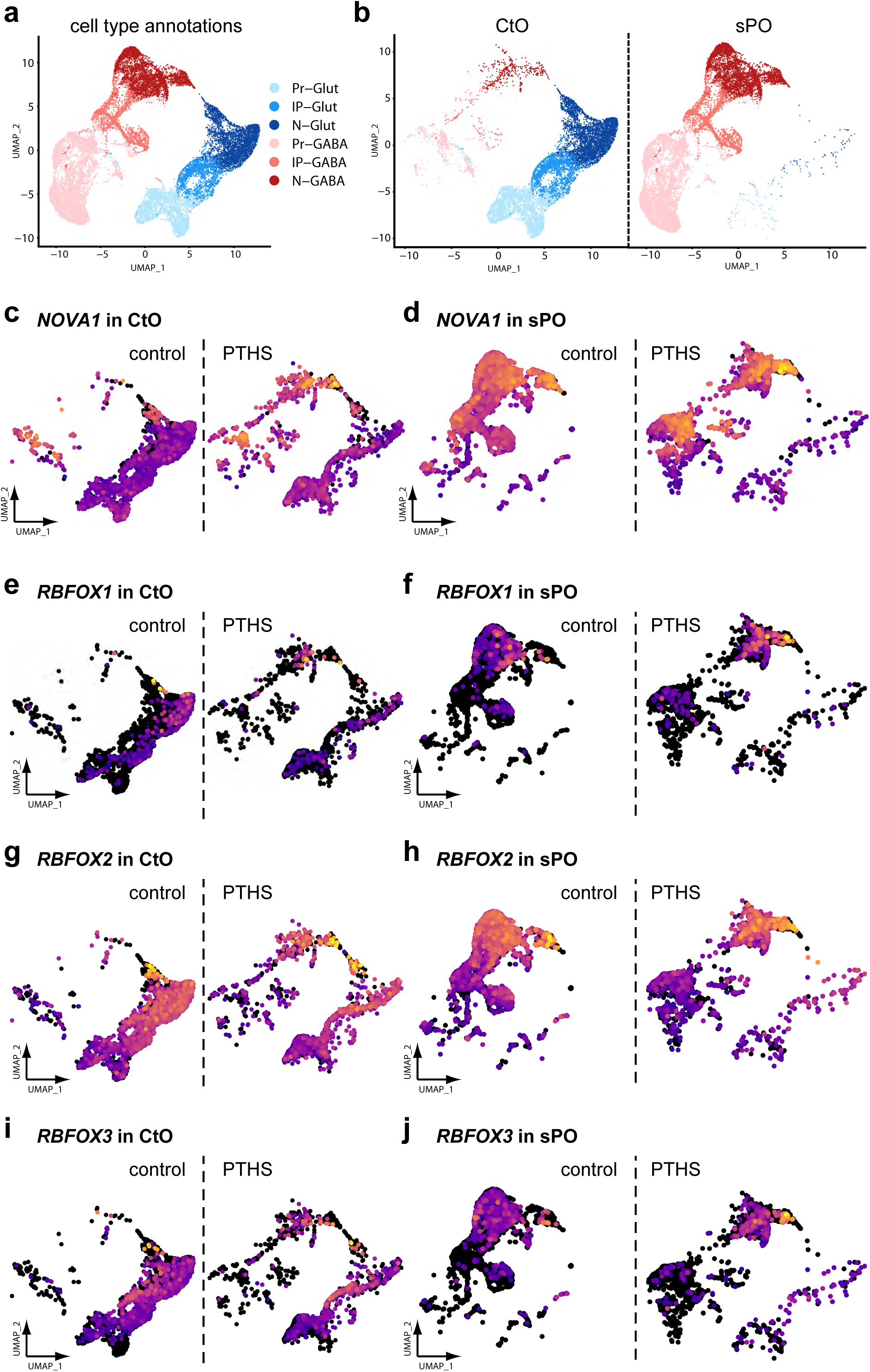
Aberrant expression of splicing regulators in PTHS cortical and subpallial organoids. **(a)** Uniform Manifold Approximation and Projection (UMAP) bidimensional reduction of scRNA-Seq profiling of cortical (CtO) and subpallial (sPO) organoids at 8 weeks in vitro, with pooled cells from 15 organoids per library. Color code represents 6 annotated subpopulations: Pr-Glut: neural progenitors in glutamatergic lineage; IP-Glut: intermediate progenitors in glutamatergic lineage; N-Glut: glutamatergic neurons; Pr-GABA: neural progenitors in inhibitory lineage; IP-GABA: intermediate progenitors in inhibitory lineage; N-GABA: neuronal population containing GABAergic interneurons. Minority subpopulations exist but were not the focus of our study. **(b)** UMAP of annotated cells in CtO and sPO organoids, separated according to organoid type, showing prevalence of excitatory neuron lineage in CtOs and preponderance of GABAergic interneurons in sPO libraries. **(c-j)** UMAPs showing expression of *NOVA1* (c,d), *RBFOX1* (e,f), *RBFOX2* (g,h), and *RBFOX3* (i,j), in control and PTHS CtOs and sPOs. Warmer colors indicate higher expression levels for each gene, while black indicates absence of detectable expression.

The fractions of cells expressing *NOVA1* are substantial in control CtO and sPO subpopulations (Fig. 2c,d) – 80.3% of CtO progenitor cells (Pr-Glut), 67.6% of CtO intermediate progenitor cells (IP-Glut), 49.1% of CtO glutamatergic neurons (N-Glut), 96.2% of sPO progenitor cells (Pr-GABA), 97.1% of sPO intermediate progenitor cells (IP-GABA), and 98.4% of sPO GABAergic neurons (N-GABA) express *NOVA1* above a threshold level of 0.5. Curiously, *NOVA1* expression remains stable along the differentiation trajectory from progenitor cells to neurons in the glutamatergic lineage of control CtOs (Fig. 2c) (mean expression of 1.77, 1.26, and 1.3 in Pr-Glut, IP-Glut, and N-Glut, respectively), whereas expression increases from progenitors to neurons in the GABAergic lineage of control sPOs (Fig. 2d) (mean expression of 5.7, 5.6, and 12.0 in Pr-GABA, IP-GABA, and N-GABA, respectively).

In contrast, the numbers of cells expressing *RBFOX1* are much lower than those expressing *NOVA1,* across all CtO and sPO subpopulations (Fig. 2e,f) – 1.56% of CtO progenitor cells (Pr-Glut), 3.21% of CtO intermediate progenitor cells (IP-Glut), 10.7% of CtO glutamatergic neurons (N-Glut), 0.6% of sPO progenitor cells (Pr-GABA), 2.0% of sPO intermediate progenitor cells (IP-GABA), and 8.8% of sPO GABAergic neurons (N-GABA) express *RBFOX1* above a threshold level of 0.5. *RBFOX1* expression increases dramatically along the differentiation trajectory from progenitor cells to neurons in both the glutamatergic and GABAergic lineages of control CtOs and sPOs, respectively (Fig. 2e,f) (mean expression of 0.04, 0.06, 0.23, 0.001, 0.03, and 0.2 in Pr-Glut, IP-Glut, N-Glut, Pr-GABA, IP-GABA, and N-GABA, respectively).

On the other hand, *RBFOX2* is expressed in a large fraction of cells in CtO and sPO subpopulations (Fig. 2g,h) – 55.4% of CtO progenitor cells (Pr-Glut), 84.9% of CtO intermediate progenitor cells (IP-Glut), 73.6% of CtO glutamatergic neurons (N-Glut), 28.9% of sPO progenitor cells (Pr-GABA), 50.3% of sPO intermediate progenitor cells (IP-GABA), and 88.5% of sPO GABAergic neurons (N-GABA) express *RBFOX2* above a threshold level of 0.5. It is very clear that *RBFOX2* expression increases markedly along the differentiation trajectory from progenitor cells to neurons in both the glutamatergic lineage of control CtOs (Fig. 2g) (mean expression of 0.98, 2.1, and 2.6 in Pr-Glut, IP-Glut, and N-Glut, respectively) and the GABAergic lineage of control sPOs (Fig. 2h) (mean expression of 0.46, 0.96, and 3.39 in Pr-GABA, IP-GABA, and N-GABA, respectively).

Although *RBFOX3* is a widely recognized neuronal marker, it is expressed in a significant fraction of progenitor cells and intermediate progenitors in CtO and sPO subpopulations (Fig. 2i,j) – 11.7% of CtO progenitor cells (Pr-Glut), 49.4% of CtO intermediate progenitor cells (IP-Glut), and 28.5% of CtO glutamatergic neurons (N-Glut), whereas the fractions of sPO cells expressing *RBFOX3* are less prominent – 0% of sPO progenitor cells (Pr-GABA), 4.4% of sPO intermediate progenitor cells (IP-GABA), and 14% of sPO GABAergic neurons (N-GABA) express *RBFOX2* above a threshold level of 0.5. Different than *RBFOX1* and *RBFOX2*, *RBFOX3* expression peaks at the intermediate progenitor stage in the glutamatergic lineage of control CtOs (Fig. 2i) (mean expression of 0.21, 1.0, and 0.53 in Pr-Glut, IP-Glut, and N-Glut, respectively) and increases marginally along the GABAergic lineage of control sPOs (Fig. 2j) (mean expression of 0.007, 0.08, and 0.25 in Pr-GABA, IP-GABA, and N-GABA, respectively).

In the comparison between PTHS and control organoids, we verified that the percentage of N-Glut neurons expressing *NOVA1* is higher in PTHS CtOs in contrast to control organoids (left panels in Fig. 3a) and that the percentage of *NOVA1+* N-GABA neurons remains practically unchanged in PTHS sPOs (right panels in Fig. 3a). Although the mean levels of *NOVA1* expression are statistically significantly different in the comparison between PTHS and control N-Glut neurons in CtOs, being higher in the patient-derived organoids, the effect size is not large (2.0 expression abundance in PTHS versus 1.3 in control; Fig. 2c). Likewise, *NOVA1* expression is statistically significantly higher in PTHS N-GABA neurons of sPOs, and the effect size is similarly not large (14.8 expression abundance in PTHS versus 12.0 in controls; Fig. 2d).

**Figure 3:**
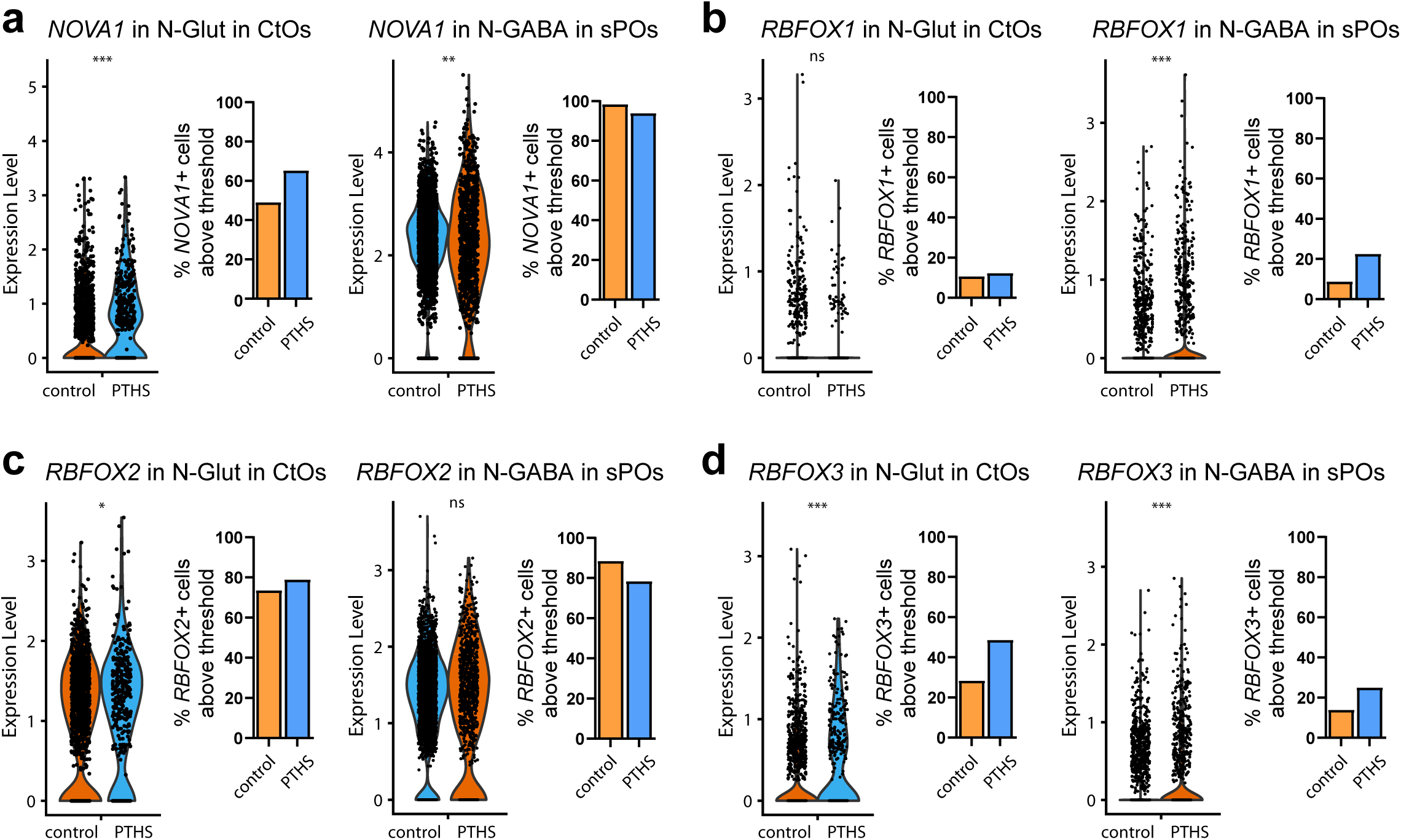
**Altered percentage of cells expressing splicing regulators in PTHS organoids**. **(a-d)** Left graphs in each group: Violin plots showing comparison of expression of *NOVA1* (a), *RBFOX1* (b), *RBFOX2* (c), and *RBFOX3* (d) between N-Glut subpopulations in the parent and PTHS CtOs (left groups) or between N-GABA subpopulations in the parent and PTHS sPOs (right groups). N = 1401 (parent) or 380 (PTHS) N-Glut cells, or N = 2661 (parent) and 988 (PTHS) N-GABA cells. Right graphs in each group: changes in percentages of neuronal subtypes in the analyzed subpopulations of CtOs and sPOs (cells expressing each gene above threshold corresponding to 0.5 log-normalized expression levels).

Similar to *NOVA1*, the percentages of N-Glut neurons expressing *RBFOX1* and *RBFOX2* are practically unchanged in PTHS CtOs in contrast to control organoids (left panels in Fig. 3b,c), the percentage of N-GABA neurons expressing *RBFOX1* is slightly higher in PTHS sPOs, and the percentage of N-GABA neurons expressing *RBFOX2* is lower in PTHS sPOs (right panels in Fig. 3b,c). Interestingly, the percentages of cells expressing RBFOX3 are higher in both CtOs and sPOs (Fig. 3d). Finally, the mean expression levels of *RBFOX1*, *RBFOX2*, or *RBFOX3* are statistically significantly dysregulated in PTHS N-Glut neurons in CtOs, with small effect sizes (0.19, 3.14, and 1.0 expression abundances in PTHS versus 0.23, 2.6, and 0.53 in control, respectively; Fig. 2e,g,i), being higher in PTHS N-GABA neurons of sPOs, with equally small effect sizes (0.74, 3.6, and 0.61 expression abundances in PTHS versus 0.2, 3.4, and 0.25 in control, respectively; Fig. 2f,h,j).

The higher expression of *NOVA1* in PTHS CtO and sPO organoids (Fig. 2c,d) as well the higher expression of *RBFOX1/2/3* in PTHS sPO organoids (Fig. 2f,h,j) contrasts with their lower expression in PTHS neurons grown in traditional 2D culture (Fig. 1d). As stated above, this difference may be due to the much wider diversity of cell types and connectivity found in the organoids. Regardless, the bulk and single-cell RNA-Seq results described here and in the previous section indicate alterations in the expression of *NOVA1* and *RBFOX1/2/3* splicing regulators in Pitt-Hopkins Syndrome. These findings prompted us to further investigate possible changes in the RNA splicing process in the patient-derived neural tissue.

### Aberrant splicing patterns in neurons derived from Pitt-Hopkins Syndrome patients

The consistent finding of altered expression of RBFOX and NOVA splicing regulators in neuronal cultures derived from PTHS patients led us to hypothesize that such dysregulated expression may have led to aberrant splicing. To investigate this, we conducted a computational analysis of the RNA-Seq data from neuronal cultures derived from 5 patient-control pairs using the software rMATS-turbo^41^, a computational tool for analyzing alternative splicing using short-read RNA-seq data. In short, rMATS-turbo works by initially using fastq input files to produce splicing graphs for each RNA-seq sample, followed by integration across samples and replicates to quantify alternative splicing events of five basic types: skipped exon (SE), alternative 5′ splice sites (A5SS), alternative 3′ splice sites (A3SS), mutually exclusive exons (MXE) and retained intron (RI). rMATS-turbo adopts the widely used PSI metric to quantify splicing, which represents the percentage of transcripts that include a specific exon or splice site, as calculated from the read counts supporting those specific exons or splice junctions, normalized by the effective lengths of the gene’s transcripts^41^. Differences between PSI values in the control and PTHS samples were then computed as ΔPSI, which can be used to infer the existence of different splicing patterns in one or another condition (for example, in PTHS neurons). Positive ΔPSI values imply that the splicing event under analysis is more prevalent in the control samples, while negative ΔPSI values indicate stronger usage in the PTHS samples.

After filtering for PSI values (averaged across biological replicates for PTHS or control conditions) smaller than 5% or greater than 95% in both samples, for a false discovery rate smaller than 5% and for |ΔPSI| greater than 5%, we could detect a significant number of splicing events with different PSI in the comparison between neurons derived from each PTHS patient against respective control, in all five categories of splicing (Supplementary Files 1-25). As an example, we could detect 3695 alternative SE events, where one exon of a gene is excluded from a transcript at a higher rate in one sample versus the other, with |ΔPSI| greater than 5% in the comparison between patient and parent #2 (Fig. 4a; Supplementary Table S2). These events were related to transcripts from 2137 different genes (Supplementary Table S2). The numbers of differential splicing events and the numbers of genes in which these events happen, in the 5 patient-control pairs, can be found in Supplementary Table S2. It should be noted that SE events are more numerous than the other types, and RI or A5SS events are the rarest (Fig. 4a). When we intersected the splicing events across the five patient-control pairs, we found 492 SE events that are common to all pairs (Fig. 4b), in 365 genes (Fig. 4c; Supplementary Table S3). The numbers of common splicing events of the other 4 types are found in Fig. 4b,c and Supplementary Table S3.

**Figure 4:**
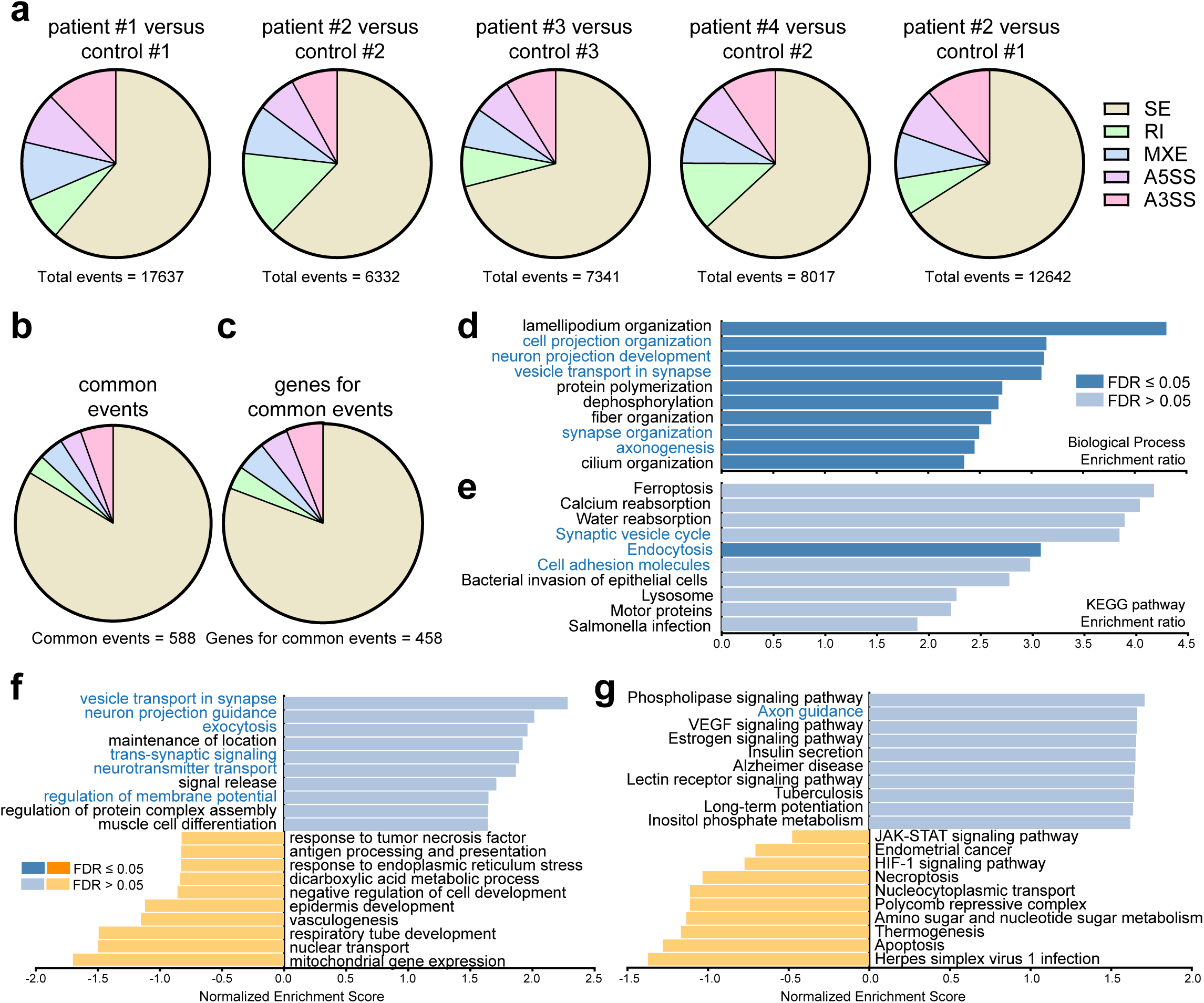
Genes exhibiting aberrant splicing patterns in PTHS are involved in synapse formation and cell-cell communication. **(a)** Pie charts representing the distribution of splicing events in 5 alternative splicing categories, in several pairs of patient and control samples. **(b)** Pie chart indicating the number of events in each alternative splicing category for the common splicing events across all patient-control pairs. **(c)** Pie chart representing the number of genes for which splicing events were detected in each alternative splicing category, in the intersection across all patient-control pairs. **(d,e)** Enrichment ratios observed in Over-Representation Analysis (ORA) of splicing events with an absolute value of DPSI larger than 0.3 in the patient-control #2 pair, subjected to either Gene Ontology – Biological Process (d) or KEGG Pathway analysis (e). Only the 10 categories with the largest enrichment ratios are shown. The color of each bar represents the false discovery rate (FDR) and the category names indicated in blue represent those linked to synapse formation and function or cell-cell communication. **(f.g)** Enrichment ratios observed in similar analyses conducted in d,e but with Gene-Set Enrichment Analysis (GSEA).

To better understand the scope of changes in PTHS neurons due to aberrant splicing patterns, we conducted Over-Representation Analysis (ORA) followed by Gene Ontology (GO) and KEGG Pathway analyses, using a list of unique genes for which rMATS-turbo detected splicing events in the comparison between each PTHS patient and a control neuronal cell line. This analysis was performed using the web-based WebGestalt tool^42^, which results in an over-representation *z-*score and enrichment *p*-value for each GO term. Interestingly, for SE events, when we analyzed the set of genes from the comparison between patient #2 and its control, we detected a significant enrichment of genes in Gene Ontology-Biological Processes (GO-BP) and KEGG-Pathway categories linked to synapse physiology, axon projection, and cell adhesion (Fig. 4d,e; Supplementary Fig. S1). GO and Pathway analyses results for other patient-control pairs yielded similar results (Supplementary Figure S1), highlighting the possibly relevant role of aberrant splicing patterns in the neural tissue cellular and molecular abnormalities that accompany PTHS.

Likewise, we conducted Gene Set Enrichment Analysis (GSEA) followed by Gene Ontology (GO) and KEGG Pathway analyses, using the same list of unique genes for which rMATS-turbo detected splicing events in the comparison between each PTHS patient and a control neuronal cell line. For these analyses, the list of genes was ordered according to the |ΔPSI| values of the corresponding splicing events from rMATS-turbo. GSEA results indicate that, among the genes at the top of the list (meaning genes with splicing events that are more dissimilarly frequent in the comparison between PTHS and control), there are genes related to synapse formation, synapse physiology, and ion transport (Fig. 4f,g; Supplementary Fig. S2). It is interesting to note that the GO and Pathway enrichment analyses described above show the presence of aberrant splicing patterns in genes related to synapse formation, which may be responsible for the lower electrical activity found in PTHS organoids^34^.

The finding of a substantial number of splicing events that seem to occur differentially between PTHS and control neuronal cell lines led us to hypothesize that the alterations found in the expression levels of *NOVA1* and *RBFOX1/2/3* in PTHS neurons are causally related to the aberrant splicing patterns observed.

### NOVA1 and RBFOX1/2/3 RNA binding sites are enriched in transcripts predicted to have aberrant alternative splicing patterns in PTHS

To further study the relationship between the aberrant splicing patterns observed in PTHS neurons and the dysregulation of expression of NOVA and RBFOX splicing regulators, we investigated the occurrence of NOVA1 and RBFOX1/2/3 binding sites in the genes detected by rMATS-turbo described in the previous section. First, we extracted the genomic sequences containing the exons involved in the SE splicing events detected by rMATS-turbo, plus additional sequences of 200 bp in the neighboring introns. The resulting collection of exon sequences flanked by adjacent introns was then analyzed for RBP (RNA binding protein) binding site enrichment using the SEA program in the MEME suite^43^. Our reasoning was that, if NOVA1 or RBFOX regulators were involved in the aberrant splicing patterns observed in PTHS, then we would expect to see an enrichment in binding sites near the splicing events relative to the occurrence of RBP binding sites in a random collection of genomic sequences. Two NOVA1 recognition motifs were analyzed^44^ and both were found to be significantly enriched in the genomic sequences near the SE events detected by rMATS-turbo (*E*-values of 9.2 × 10^-8^ and 3.0 × 10^-8^; Fisher’s exact test; Fig. 5a), with a lower *E*-value than those calculated for a set of genes known to be regulated by NOVA1 (positive control; *E*-values of 2.9 × 10^-1^ and 1.6 × 10^-6^, respectively; Fig. 5a; Supplementary File S4). The same type of analysis was conducted for binding site motifs recognized by RBFOX1 and RBFOX2/3 splicing regulators^44^, indicating that both are significantly enriched in the genomic sequences near the SE events detected by rMATS-turbo (*E*-values of 5.8 × 10^-4^ and 4.1 × 10^-7^, respectively; Fisher’s exact test; Fig. 5a); although these values are larger than those calculated for a set of genes known to be regulated by RBFOX1/2/3 (positive control; *E*-values of 8.4 × 10^-8^ and 9.2 × 10^-16^, respectively; Fig. 5a; Supplementary File S4), the *E*-values are still much smaller than when the enrichment analysis was performed by comparing two randomly chosen sets of genomic sequences (Fig. 5a).

**Figure 5:**
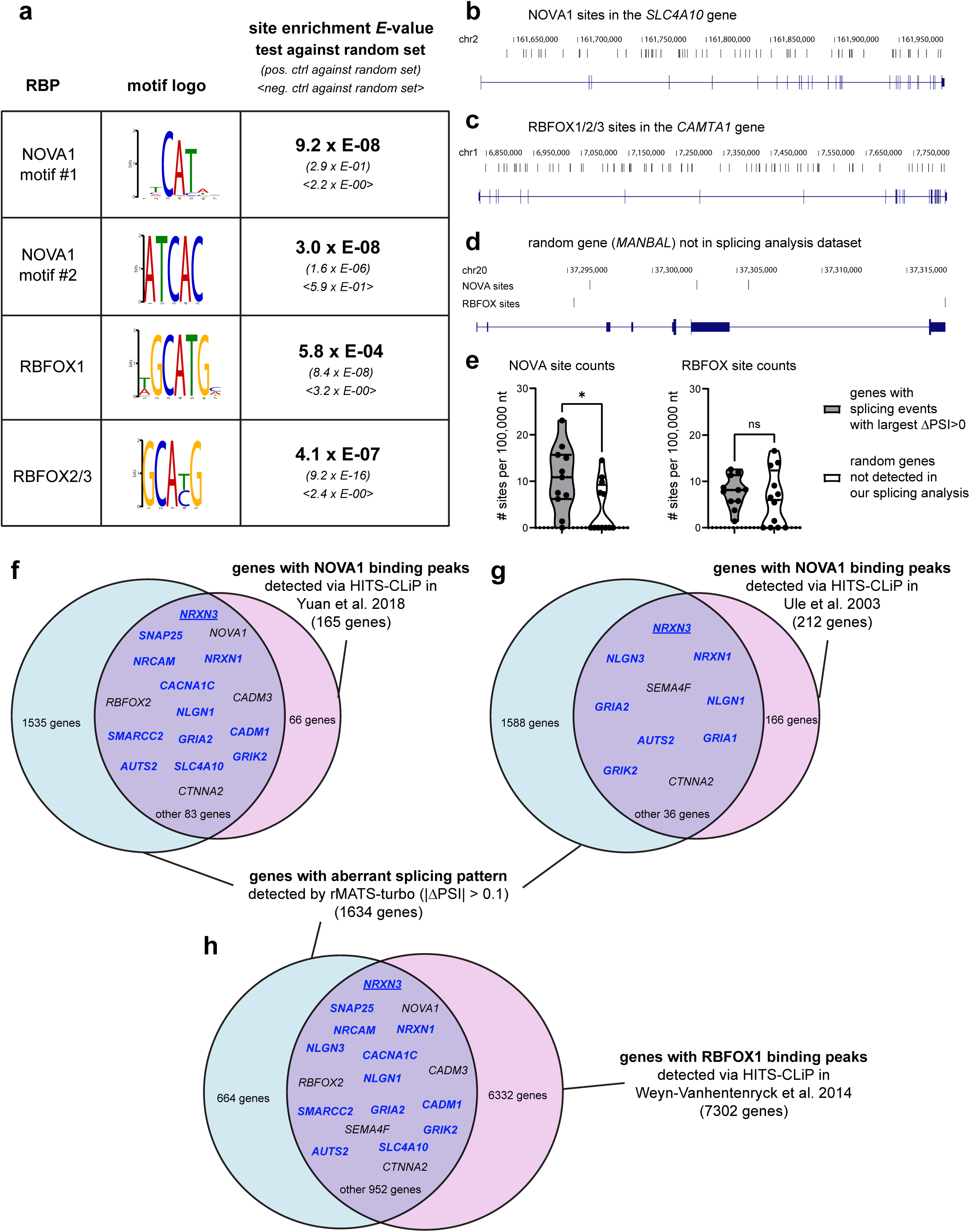
Enrichment of binding sites for splicing regulators NOVA1 and RBFOX1/2/3 in genes with altered splicing patterns in PTHS. **(a)** Results from site enrichment analysis conducted on the sequences of skipped exons and neighboring introns for splicing events detected by rMATS-turbo in PTHS neurons, for RNA-binding proteins (RBP) NOVA1 and RBFOX1/2/3, with two motifs for NOVA1, one motif for RBFOX1 and one common motif for RBFOX2 and RBFOX3 (motif logos in the middle column). The statistical results from Fisher’s exact test (*E*-value) in the comparison between the test dataset (sequences from splicing events in PTHS) and genomic sequences from a random set of genes are found in boldface text. For contrast, the *E*-values for the comparison between a positive control dataset (sequences of genes known to be regulated by the corresponding RBP) and the random control dataset are given in parentheses, and the *E*-values for the comparison between a negative control dataset (sequences of genes not known to be regulated by the corresponding RBP) and the random control dataset are given between <> symbols. **(b,c)** Examples of genomic sequence structure for two genes in our aberrant splicing dataset detected by rMATS-turbo, including the *SLC4A10* gene, shown to be enriched for NOVA1 binding sites (b) and the *CAMTA1* gene, shown to be enriched for RBFOX1/2/3 binding sites (c). The exons are shown as blue rectangles, connected by lines representing the introns. The binding sites are shown as black bars above the gene representation, and the chromosome numbers and genomic coordinate are also displayed. **(d)** Representation of NOVA1 and RBFOX1/2/3 binding sites in a random gene (MANBAL) not present in the aberrant splicing event list from PTHS neurons. **(e)** Violin plots representing the statistical comparison of the numbers of NOVA1 (left graph) and RBFOX1/2/3 (right graph) sites, in 100,000 nt, in the genes with the highest positive values of ΔPSI from the rMATS-turbo analysis of PTHS neurons (n=11 genes) compared with a random set of genes not present in the splicing event list in PTHS (n=12 genes). The median/interquartile range are shown as horizontal bars inside the violins. Welch’s unpaired t-test assuming unequal variances; * *p*<0.05; ns, not statistically significantly different. **(f-h)** Venn diagrams representing the comparison between the number of genes with differential splicing events in PTHS neurons, as detected by rMATS-turbo in RNA-Seq data (absolute values of ΔPSI larger than 0.1), and the number of genes whose mRNA was shown in the literature to bind NOVA1 (f; ref ^46^), NOVA1 (g; ref ^63^), and RBFOX1 (h; ref ^1^), in HITS-CliP experiments. The numbers of genes in each sector are shown inside the Venn diagrams, and some genes involved in neurodevelopment are highlighted in each intersection. Genes highlighted in blue are those present in the SFARI database of autism-related genes^78^.

Next, we conducted an experiment to detect putative RBP binding sites across the entire genomic sequence of the genes whose splicing patterns were found to be dysregulated by rMATS-turbo. This analysis was performed using the RNA-binding protein motif detection software FIMO in the MEME suite^45^. Examples of genes shown to have enrichment of NOVA1 and RBFOX1/2/3 sites are shown in Fig. 5b,c, in comparison with the lower abundance of binding motifs in genes in the random set (negative control; Fig. 5d). We observed a difference in site count which is statistically significant for NOVA1 (Fig. 5e) when considering the genes for the top splicing pattern alterations, as measured by the absolute value of ΔPSI (larger than 30%).

These enrichment analyses suggest that NOVA1 and RBFOX splicing regulators may be involved in dictating the aberrant splicing patterns observed in PTHS neurons. Direct evidence for this relationship was further pursued by the analysis of published CLiP-Seq (HITS-CLiP) experimental datasets, from studies that performed transcriptome-wide determination of NOVA and RBFOX binding sites^1,11,46^. Considering the 1634 genes harboring aberrant SE splicing events with absolute ΔPSI values larger than 0.1 (larger splicing usage either in the parent or the patient), there is a substantial overlap with the hundreds of genes whose transcripts were shown to be directly bound to NOVA1 (Fig. 5f,g; Supplementary File S5), including genes coding for synapse organization and neuronal physiology. Likewise, there is a very substantial overlap with the thousands of genes whose transcripts have been shown to be directly bound to RBFOX1/2/3 splicing regulators (Fig. 5h; Supplementary File S5).

In combination, these results suggest that the altered splicing patterns detected in the transcriptomes of PTHS neurons may be caused by the dysregulation of NOVA1 and RBFOX1/2/3 splicing regulators.

### The Neurexin-3 gene (*NRXN3*) exhibits an exon skipping event that leads to a transcript encoding a soluble protein isoform

Considering that the list of skipped exon (SE) splicing events detected by rMATS-turbo is longer than the lists for the other types of differential splicing events (Fig. 4a), we decided to further investigate this category to better understand the types of events and genes affected by aberrant splicing patterns. GO and Pathway analyses conducted on the list of genes for which rMATS-turbo detected splicing events revealed the presence of genes in enriched categories related to synapse formation and physiology, axonal projection, and cell adhesion (Fig. 4d-g). Curiously, we detected a significant number of splicing events that occur in genes in the Neurexin family, which are traditionally regarded as critical components of synapse physiology^47^.

In the list of SE splicing events in the comparison between patient-control pairs #1, #2, and #3, we could detect 6, 12, and 1 events, respectively, in genes in the Neurexin family (*NRXN1, NRXN2,* and *NRXN3*) (Supplementary File S2). Because some qPCR experiments described below were conducted comparing patient #2 with the parent #1 control (due to non-availability of parent #2 samples), we also conducted the parent #1/patient #2 comparison using rMATS-turbo, which yielded 9 differential SE events linked to *NRXN1/2/3* genes. Finally, in the comparison between parent #2 and patient #4, for whom there is no associated parent in our cohort, we could also detect 2 SE events linked to *NRXN2* and *NRXN3* genes (Supplementary File S2).

The numerous events related to *NRXN* genes are further highlighted by the presence of 2 events associated with *NRNX2* and *NRXN3* among the top 20 SE splicing events with positive ΔPSI (higher usage in control samples) in the patient #2-parent #2 comparison (Supplementary File S2). The event in the *NRXN2* gene has a ΔPSI equal to 0.681 (Fig. 6a,b), with a higher splicing event usage (ranging from 0.797 to 0.86) in the parent #2 biological replicate libraries, while the PSI values range from 0.117 to 0.196 in the patient #2 samples (Supplementary File S2). It is interesting to note that the exon preferentially included in the transcripts of the *NRXN2* gene in the control sample (SE-NRXN2-i splicing event) is very short (24 nt; Fig. 6b,e) and is equivalent to exon 7 in the *NRXN2* Ensembl transcript ENST00000409571.6. In the PTHS patient, exon 7 is preferentially excluded (SE-NRXN2-e splicing event (Fig. 6b). Interestingly, there exists a 45-bp annotated exon in the same genomic region for the canonical *NRNX2* Ensembl transcript (accession ENST00000265459.11) (Fig. 6c). The only difference between these two annotated transcripts is the 21-bp difference in this exon. The exon skipping event does not lead to a change in the coding sequence reading frame and results in a loss of a few amino acid residues relative to the protein encoded by the canonical transcript (Fig. 6c); therefore, we decided not to pursue this splicing event further.

**Figure 6:**
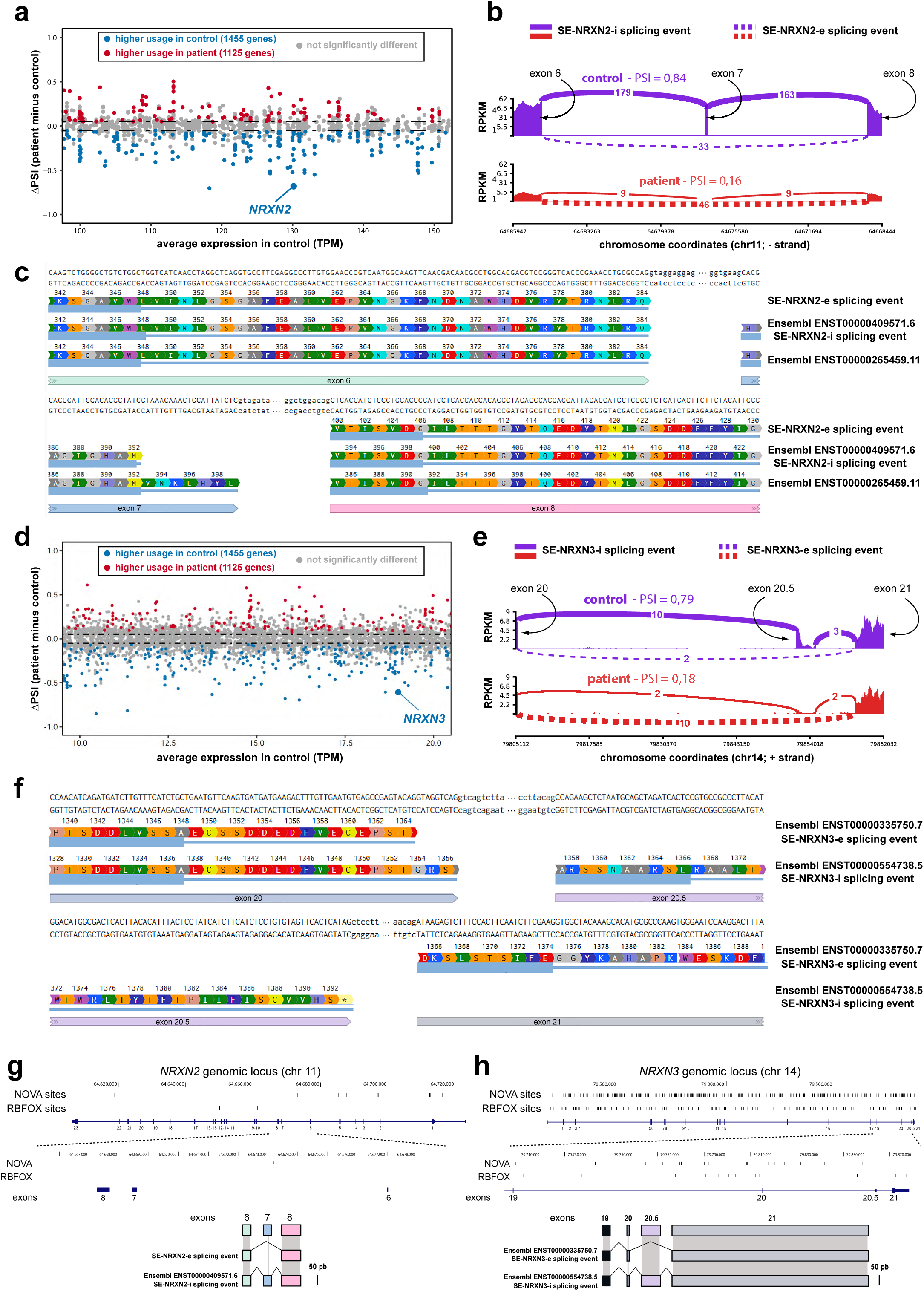
Aberrant splicing in Neurexin genes in PTHS. **(a)** MA plot representing the differential splicing event prevalence, in the comparison between the PTHS patient-derived neurons and control neurons cultivated in vitro. The *x*-axis represents the expression abundance for each gene (represented as dots), in transcripts per million (TPM), while the *y*-axis shows the difference between the PSI (percent spliced in) for the patient cells and the control neurons (ΔPSI). Each dot represents a gene. Only a fraction of the genes is shown, to highlight the region of the graph where we observed a splicing event in the *NRXN2* gene chosen for further analysis. Blue dots indicate splicing events with a statistically significantly higher usage in control neurons, involving the number of genes displayed in parenthesis. The red dots indicate splicing events with higher usage in patient neurons. Gray dots are splicing events with a non-statistically significant difference in usage between patient and control cells. **(b)** Sashimi plots representing the splicing patterns for the NRXN2 splicing event highlighted in (a), depicting the location of exons 6, 7, and 8 in chromosome 11 (genomic coordinates below the plots). The vertical bars indicate the coverage of RNA-Seq reads in each exon or supporting the existence of each splice junction, in a *y*-axis scale shown as RPKM (reads per kilobase per million total reads). The cusp lines connecting the exons above or below the vertical bars indicate reads that support the existence of each corresponding exon-exon junction, with full lines representing junctions that include SE (skipped exon) 7, while dashed lines display functions that exclude said exon. The cusp thickness is proportional to the number of reads supporting the existence of each splice junction. The Sashimi plot on top (purple) represents the splicing pattern in control neurons, and the plot at the bottom (red) indicates the pattern in PTHS neurons, with the PSI values indicated above each graph. **(c)** Schematic representation of a fraction of the genomic sequence of NRXN2 gene, showing exons 6, 7, and 8. Three transcripts are shown: one represents the transcript that excludes exon 7 (SE-NRXN2-e), as detected by our rMATS-turbo analysis of PTHS neurons; the second represents the transcript including exon 7 (SE-NRNX2-i), which also corresponds to a transcript in the Ensembl database (ENST entry number shown); the transcript at the bottom is another Ensembl transcript that includes a slightly longer version of exon 7. The amino acid translations for each transcript are shown below the DNA sequences. **(d)** Similar to (a), but for an event involving the *NRXN3* gene, chosen for further analyses. **(e)** Similar to (b), but for the event in (d), involving exons 20 and 21 of the canonical transcript in the Ensembl database, and an additional exon not annotated in the canonical transcript (exon 20.5). **(f)** Similar to (c), showing two transcripts of the *NRXN3* gene, either including (SE-NRXN3-i) or excluding (SE-NRXN3-e) exon 20.5. **(g,h)** Genomic sequence structure for NRNX2 (g) and NRNX3 (h), showing the location of NOVA1 and RBFOX1/2/3 binding sites. The exons are shown as blue rectangles (exon numbers below the gene structure), connected by lines representing the introns. The binding sites are shown as black bars above the gene representation, and the chromosome numbers and genomic coordinate are also displayed. The region containing the exons involved in the SE events are blown up below. The models at the bottom schematically depict the exons (to scale) and the splicing patterns in the SE-NRXN2/3-i and SE-NRXN2/3-e transcripts.

On the other hand, the SE event in the *NRXN3* gene has a ΔPSI equal to 0.608 (Fig. 6d,e), with splicing event inclusion levels ranging from 0.579 to 1.00 in the parent #2 biological replicate libraries, while the PSI values range from 0.167 to 0.185 in patient #2 samples (Supplementary File S2). For ease of reference, we will name the SE event that includes the exon as SE-NRXN3-i, while the splicing configuration that does not include the exon will be called SE-NRXN3-e. Moreover, we chose to call the included exon as E20.5, thus named because it is included between exons 20 and 21 of the canonical *NRXN3* Ensembl transcript ENST00000335750.7 (this exon is also equivalent to exon 19 in the *NRXN3* Ensembl transcript ENST00000554738.5). The exon that is preferentially included in the control samples has a size of 679 nt (Fig. 6e,f). This sequence is found in only 2 out of 14 *NRNX3* Ensembl transcripts with recognized coding sequences (ENST00000554738.5 and ENST00000555387.1); on the other hand, this is not an annotated exon in the canonical *NRXN3* Ensembl transcript (ENST00000335750.7) (Fig. 6f). Additionally, this included exon has been described before in the context of *NRXN3* transcripts expressed in post-mortem human cerebral cortex samples of individuals harboring a SNP near a splicing site in the *NRXN3* gene (rs8019381)^48^. As will be described in detail below, this included exon creates a premature stop codon and the resulting protein truncating transcript variant (PTV) leads to a significantly shorter NRNX3 protein isoform.

Interestingly, when we analyzed the sequences of *NRXN2* and *NRXN3* genes using the RNA binding protein binding site motif detection software FIMO^45^ (MEME suite), we could detect 4 and 223 NOVA1 binding sites in *NRXN2* and *NRNX3* (Fig. 6g,h), respectively. Of these, 1 site falls into adjacent introns and none falls inside the skipped exon for the *NRXN2* event (Fig. 6g), whereas 9 sites fall inside the flanking introns and none falls in the skipped exon for the *NRXN3* splicing event (Fig. 6h). Likewise, we could detect 13 and 156 RBFOX1/2/3 binding sites in the *NRXN2* and *NRNX3* genes (Fig. 6g,h), respectively. Of these, none falls into neighboring introns or inside the skipped exon for the *NRXN2* event (Fig. 6g), while one falls inside the flanking introns and none falls inside the skipped exon for the *NRXN3* splicing event (Fig. 6h).

In combination, these results argue in favor of the *NRNX2* and *NRXN3* events under analysis are controlled by NOVA and RBFOX splicing regulators. To confirm the differential splicing in the *NRXN3* gene that creates the SE events mentioned in the previous paragraphs, we decided to conduct reverse-transcription-coupled real-time quantitative PCR (RT-qPCR) as well as standard PCR amplification, using gene-specific primers able to discriminate between those splicing events. To detect the SE-NRXN3-i variant, we placed the forward oligo (p5) in exon 19 of the canonical transcript and the reverse oligo (p4) in the splice junction between exons 20 and 20.5 (Fig. 7a, right panel). To detect the SE-NRNX3-e variant, we used another forward oligo (p1) in exon 19, and the reverse oligo (p3) was placed in the splice junction between exons 20 and 21 of the canonical *NRXN3* Ensembl transcript ENST00000335750.7. To detect both variants (and the vast majority of transcripts, according to the *NRXN3* Ensembl entry), we used forward and reverse oligos (p1 and p2) in exons 19 and 20, respectively (Fig. 7a, left panel).

**Figure 7:**
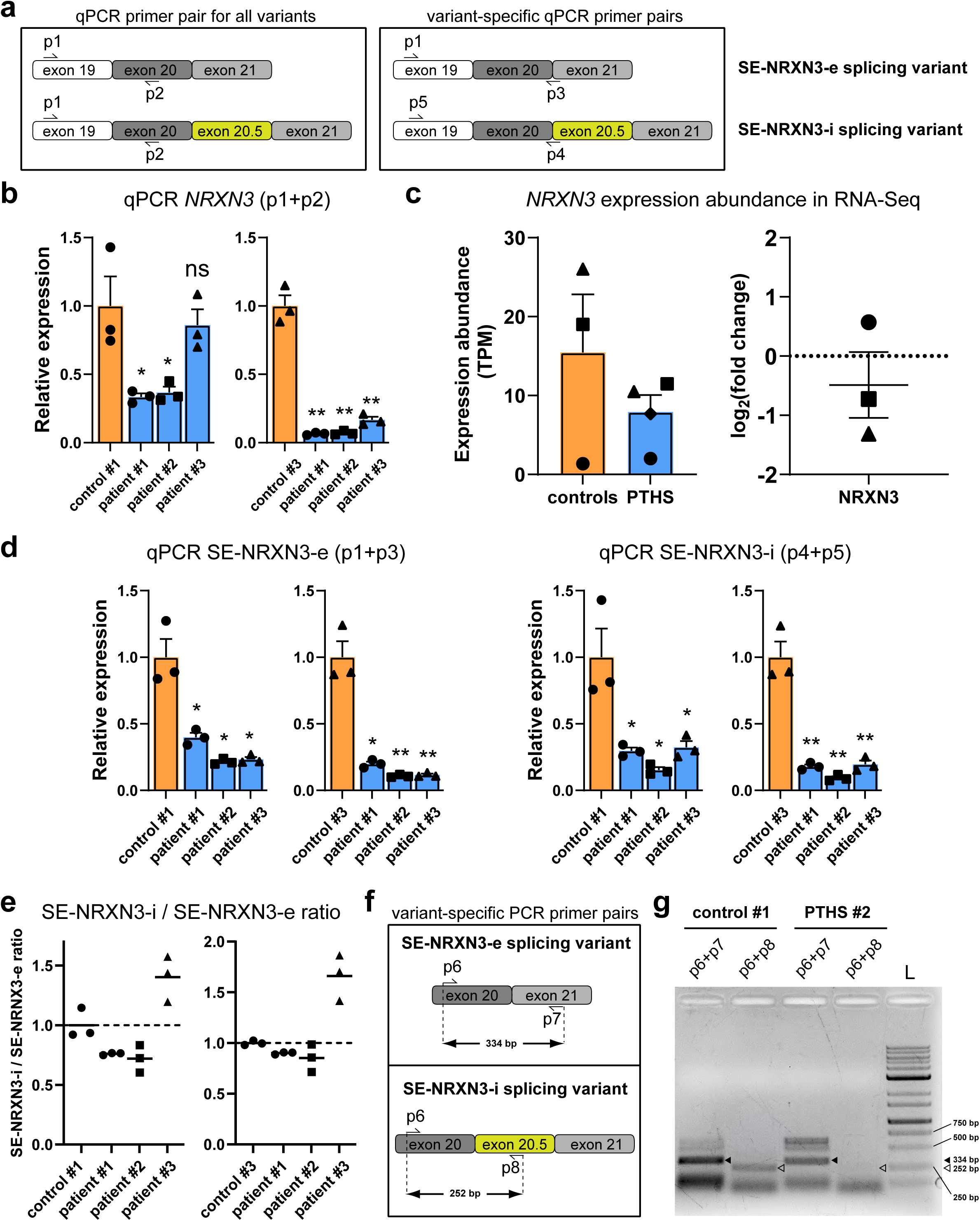
Changes in the abundances of transcript variants via exon skipping at the Neurexin-3 gene *NRXN3* in PTHS neurons in vitro. **(a)** Schematic representation of the location of PCR primers in exons of the *NRXN3* gene, used in real-time qPCR experiments. Some oligos were designed to detect all transcript variants (left), because they detect a combination of exons present in both the transcript including (SE-NRXN3-i) and excluding (SE-NRXN3-e) exon 20.5. Other PCR oligos (right) were designed as variant-specific primers, because they recognize exon-exon splice junctions typical of either the SE-NRXN3-I or SE-NRXN3-e transcripts. **(b)** Real-time qPCR results showing the expression of *NRXN3* with the combination of oligos p1 and p2, which recognize both transcript variants. Each graph compares the expression in three patient lines (blue bars) and control line #1 (orange bar on the left graph) or control line #3 (orange bar on the right graph). Error bars are S.E.M. Each symbol represents a patient line, according to Supplementary Table S6. The expression is each graph is normalized to 1 in the control line, and the statistical comparisons (Welch’s unpaired t-test assuming unequal variances) are between each patient line and the control shown in each graph. * *p*<0.05; ** *p*<0.01; ns, non-statistically significantly different. N=3 biological replicates per group. **(c)** Left: Bar plots representing the expression abundances of NRXN3 in control (orange) and PTHS (blue) neuronal cell lines, as judged by transcripts per million (TPM) in RNA-Seq data. Each symbol represents a patient or control. The control group has 3 subjects, while the PTHS group has 4 subjects; in three cases, the PTHS patient lines are matched with the corresponding parental control lines, and a fourth PTHS neuronal line is not matched to controls. Error bars are S.E.M. Notice the tendency for lower mean expression in the PTHS group, but the difference fell below the statistical significance threshold level, due to the low expression values in patient-parent pair #1 (circle symbols). Right: Direct comparison between the TPM expression abundances in three patient-parent pairs, represented as fold-change in the log_2_ scale. Error bars are S.E.M. **(d)** qPCR results with variant-specific oligos to detect the SE-NRXN3-e transcript (p1 and p3) or the SE-NRXN3-i (p4 and p5). The cell lines and comparisons are the same as in (c). Welch’s unpaired t-test assuming unequal variances; * *p*<0.05; ** *p*<0.01. **(e)** Ratio between the relative expression of SE-NRXN3-i and SE-NRXN3-e transcript abundances, produced by contrasting the RQ (relative quantitation) levels with oligos p4+p5 with those obtained with primers p1+p3. In each graph, the mean ratio was normalized to 1 in either control #1 (left) or control #3 (right) neuronal cell lines. N=3 independent biological replicates. Notice the change in the balance between the two transcript variants in two patient lines, with a reduction in the abundance of the transcript containing exon 20.5, as indicated by the mean ratio below 1 (dashed line). (f) Schematic representation of the location of primers for regular PCR amplification of specific *NRXN3* gene transcripts: oligos p6+p7 were designed to amplify just the SE-NRNX3-e variant, as p7 recognizes exon 21; oligos p6+p8 on the other hand were designed to amplify just the SE-NRNX3-i variant, as p8 recognizes the skipped exon 20.5 (yellow). (g) Image of the results of gel electrophoresis showing amplification of several bands with the p6+p7 and p6+p8 primer combinations, with samples from patient line #2 and control line #1. L: Kasvi 1 kb DNA ladder. The open triangle indicates the 252-bp band expected from the SE-NRXN3-i transcript, while the filled triangle shows the 334-bp band expected from the SE-NRXN3-e transcript. Notice the almost complete absence of the band from the SE-NRXN3-i transcript in PTHS samples.

The mean overall levels of *NRXN3* expression are lower in PTHS individuals, as judged by qPCR with the oligos to globally detect the gene’s transcripts (Fig. 7b), which is in keeping with the lower transcriptomic expression found in samples from two PTHS subjects in RNA sequencing experiments (Fig. 7c)^34^. Additionally, the combinations of oligos to detect the SE-NRXN3-i and SE-NRXN3-e splicing variants showed that both have lower expression in PTHS samples (Fig. 7d). Importantly, we calculated the ratio between the expression abundances detected via qPCR for the SE-NRXN3-i versus the SE-NRXN3-e events; compared with the ratios found in patients #1 and #2, controls #1 and #3 have a higher ratio, indicating that the SE-NRXN3-i event is proportionally more abundant than the SE-NRXN3-e event in the control samples in the comparison with the ratio found in those PTHS patient neurons (Fig. 7e).

Likewise, these numbers allow us to conclude that the difference in abundances for NRXN3-i and SE-NRXN3-e events is less prominent in patients #1 and #2 than in parents #1 and #3. Curiously, patient #3 also presents an imbalance between both variants, in this case in the opposite direction, which also represents an aberrant splicing pattern with potential functional consequences.

To further investigate the SE-NRXN3-i and SE-NRXN3-e splicing events, we used oligos for regular PCR amplification, designed to discriminate between variants based on amplicon size. One pair of primers was placed in exons 20 and 21 (Fig. 7f, top panel); the expected band size for the PCR performed on the SE-NRXN3-e event with this primer pair (p6 and p7) is 334 bp (Fig. 7f), although we could rule out the possibility that other band sizes would exist for other configurations of spliced sites involving exons 20 and 21. Another primer pair (p6 and p8) was placed in exons 20 and 20.5, which is expected to lead to band of size 252 bp only from the SE-NRXN3-i event (Fig. 7f, bottom panel). The PCR results confirm the expected amplicons for these primer pair/splicing event combinations (Fig. 7g), providing experimental evidence that confirms the in silico analyses executed by rMATS-turbo on our PTHS neuron RNA-Seq data, which predicted the existence of the SE-NRXN3-i and SE-NRXN3-e splicing variants.

The PCR results showed that we could amplify most of the expected bands that would support the existence of alternative splicing of exon 20.5. However, other bands were also amplified (Fig. 7g), therefore, we decided to clone the amplicons in bulk in a PCR cloning vector, followed by Sanger sequencing of several clones. With the primer pair positioned in exons 20 and 21, we detected amplicons with an exon configuration that matches the expected joining of exons 20 and 21 without the intervening exon 20.5 in both the control and PTHS samples (Supplementary Fig. S3a-c), and the percentage of clones supporting the existence of this exon configuration is larger in PTHS than in control samples (Supplementary Fig. S3c), which concurs with the results from rMATS-turbo showing that transcripts with SE-NRXN3-e events are more prevalent than SE-NRXN3-i in PTHS samples. Importantly, the combination of oligos pairing with exons 20 and 20.5, which is expected to amplify only the SE-NRXN3-i event-related amplicons, resulted in sequences that match this expected exon configuration in the control samples (Supplementary Fig. S4a-c); moreover, we could only find clones matching this exon configuration in control samples (Supplementary Fig. S3c), which concurs with the positive ΔPSI for the SE-NRXN3-i event in the controls (Fig. 6e). It is also curious to note that, although this type of experiment is not quantitative, the intensity of the 334-bp band with oligos in exons 20 and 21 as well as the 252-bp band with oligos in exons 20 and 20.5 are stronger in the control samples (Fig. 7g), which is in keeping with the lower global expression levels for the *NRXN3* gene in PTHS samples revealed by the qPCR experiments (Fig. 7b,c).

Although other bands were detected in these PCR experiments, some larger than the predicted size for the corresponding amplicons, they did not yield sequence reads in the Sanger sequencing experiments performed later. The significance of these bands remains to be investigated. In combination, the PCR results presented above confirm the existence of a transcript containing an SE exon in the *NRXN3* gene (SE-NRXN3-i event) identified in silico in our RNA-Seq data and that this splicing event has a lower usage in PTHS neurons, as compared to the transcript without exon 20.5 (SE-NRXN3-e event).

### The PTHS-depleted alternatively spliced *NRXN3* transcript leads to a prematurely stopped soluble Neurexin-3 protein isoform

Next, we investigated the Neurexin-3 protein isoforms coded for by transcripts containing the SE-NRXN3-i and SE-NRXN3-e events. The alternative *NRXN3* Ensembl transcript ENST00000554738.5, containing the splicing configuration with exon 20.5 (SE-NRXN3-i event), which is depleted in PTHS, results in a protein isoform with 6 laminin G domains and 3 EGF-like domains (Fig. 8a), as judged by protein domain prediction with the InterPro tool^49^.

**Figure 8:**
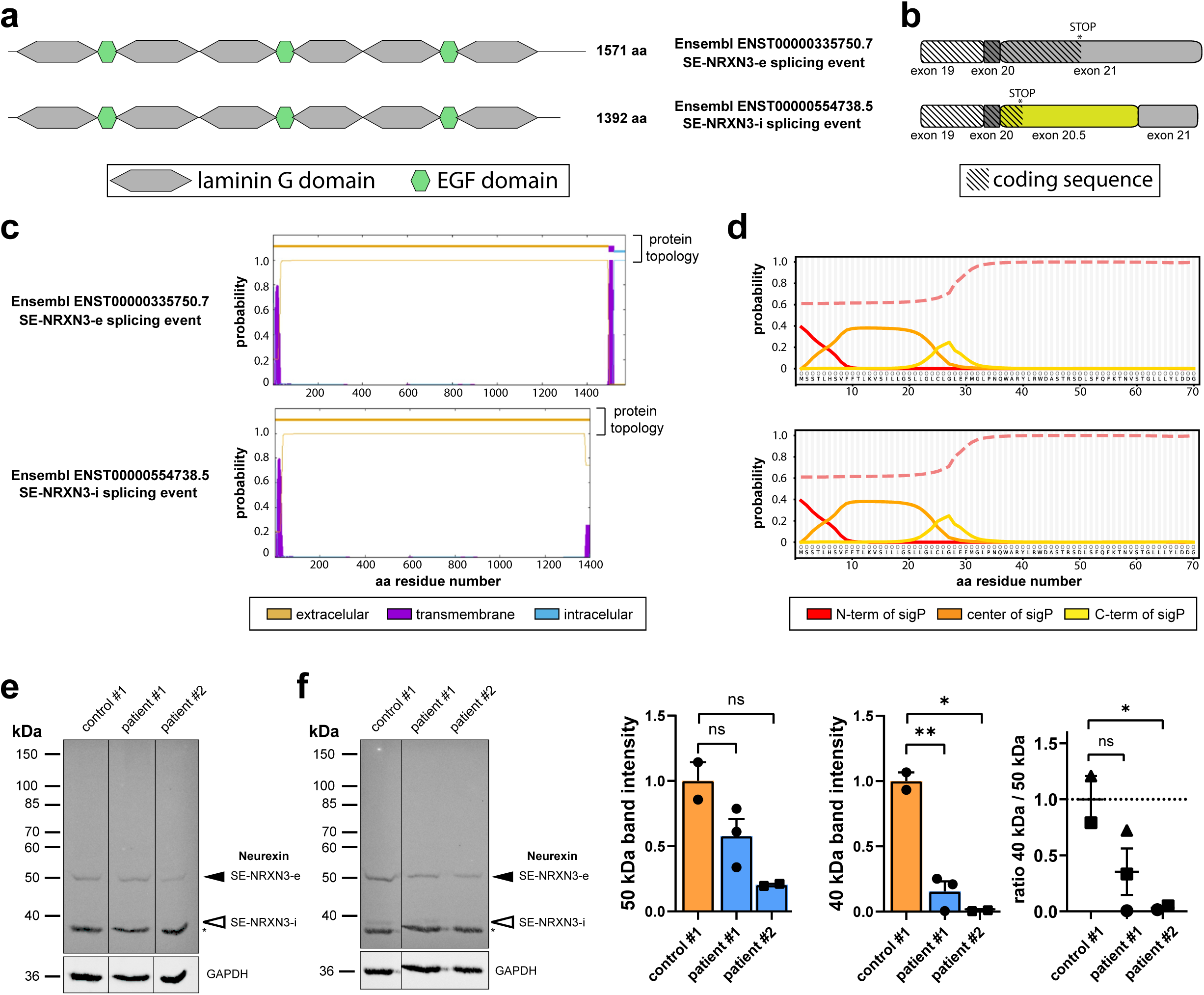
Evidence that the transcript variants of *NRXN3* encode transmembrane and secreted Neurexin-3 protein isoforms. **(a)** Location of protein domains laminin G and EGF in the Neurexin-3 protein isoforms encoded by the SE-NRNX3-e and SE-NRNX3-i variants, as detected by InterProScan, showing the shorter length of the protein predicted to be produced from the variant included exon 20.5, due to the presence of a premature stop codon**. (b)** Schematic representation showing the coding sequences in transcripts SE-NRNX3-e and SE-NRNX3-i (hatched pattern), showing a detail of the premature stop codon introduced when exon 20.5 is spliced into the transcript. Exon sizes are shown to scale, except for exon 21 in the bottom transcript, which is shown partially. **(c)** Prediction of protein topology using the DeepTMHMM tool on the proteins encoded by the SE-NRNX3-e (top) and SE-NRNX3-i (bottom) transcripts. The protein topology in each case is shown above the graph, color-coded according to the prediction of extracellular, intracellular, or transmembrane domains. The *y*-axis in the graph represents the probability of each portion of the protein being extracellular (orange), transmembrane (purple), or intracellular (blue). **(d)** Prediction of signal peptide and probability of secretion with the SignalP tool, on the first 70 amino acid residues in the protein sequences predicted from the SE-NRNX3-e and SE-NRNX3-i transcripts. Each line indicates the probability of the existence of a signal peptide (sigP) having features of N-terminal, central, or C-terminal portions of the signal peptide, while the dashed line is the remainder of the probability. **(e)** Western Blotting analysis of crude extracts from neuronal cultures in vitro, from patients #1 and #2, along with control #1. Three Neurexin bands are detected, including a ∼50 kDa band (filled arrowhead) corresponding to the predicted molecular weight of Neurexin-β, as well as ∼40 kDa bands (open arrowheads) with the predicted molecular weight of Neurexin-β lacking transmembrane domains. The endogenous control is GAPDH in all samples. The molecular weight markers shown on the left are from the PageRuler Unstained Protein Ladder (Thermo Fisher Scientific). **(f)** Similar to **e**, but with samples from cortical organoids. The graphs on the right show the quantitation of band intensity for the ∼50 kDa and ∼40 kDa bands, as well as the ratio between both bands, normalized against the control #1 sample. Notice the nearly absent expression of one of the ∼40 kDa Neurexin-β bands, whose intensity decreases more prominently than the ∼50 kDa band.

Comparatively, the canonical *NRXN3* Ensembl transcript ENST00000335750.7 encodes a protein with the same number of protein domains, but shorter C-terminus (Fig. 7A). The shorter protein encoded when exon 20.5 is included results from a premature stop codon in this exon (Fig. 8b). Because Neurexin proteins are transmembrane (TM) cellular components involved in synapse organization^47^, with the TM domain positioned near the C-terminus, we reasoned that the shorter Neurexin isoform encoded by the transcript including exon 20.5 might be lacking part of or all the TM domain. Therefore, we conducted a prediction of TM domains using the DeepTMHMM tool^50^, which showed that the shorter isoform completely lacks the TM domain and may represent a soluble form of Neurexin (Fig. 8c). Because the balance between a transmembrane and a secreted soluble form of Neurexin may have important functional implications, we checked whether the protein isoforms from the canonical and the alternative *NRNX3* transcripts are predicted to be secreted, using the SignalP (v 6.0) tool^51^. Both the complete Neurexin isoform including exon 20.5 and the shorter isoform encoded by the transcript including exon 20.5 are predicted to be secreted, because they contain a signal peptide (Fig. 8d).

To further investigate the Neurexin protein isoforms in neuronal cultures from PTHS patients and controls, we performed Western Blotting with samples harvested from neuronal cultures differentiated over the course of 3 weeks in vitro. Analysis of samples from two PTHS subjects (patients #1 and #2) and a control cell line (control #1) indicates that several protein variants are detected across all samples (Fig. 8e), including a band of molecular weight ∼50 kDa (filled arrowhead in Fig. 8e), which likely corresponds to Neurexin-3β^47^, the smaller isoform translated from a shorter mRNA transcribed from an internal alternative promoter in the *NRXN3* gene (Supplementary Table S9). This isoform is predicted to have a transmembrane domain and therefore be derived from translation of the transcript not containing exon 20.5 (SE-NRXN3-e splicing event). We could also detect bands of ∼40 kDa (open arrowheads in Fig. 8e), which is the predicted molecular weight of Neurexin-3β lacking a transmembrane domain (Supplementary Table S9), resulting from the inclusion of exon 20.5 (SE-NRXN3-i splicing event). We did not detect a band of ∼170 kDa, which would correspond to the longest Neurexin-3 isoform, called Neurexin-3α^47^, neither could we verify the presence of a band of ∼150 kDa, corresponding to the shorter version of Neurexin-3α lacking a transmembrane domain; these isoforms are predicted to be encoded by transcripts lacking or containing exon 20.5, respectively. We verified that the larger ∼50 kDa isoform is weaker in PTHS patient #2 sample as compared to the control sample (control #1) (Fig. 8e), in keeping with the observation that the overall *NRXN3* expression levels are reduced in PTHS neurons from patient #2 in qPCR and RNA-Seq experiments (Fig. 7b,c).

Importantly, we performed Western Blotting analysis of PTHS and control cortical organoids (see Methods for details) and verified that both the larger ∼50 kDa isoform and one of the ∼40 kDa bands are weaker in PTHS patients #1 and #2 samples as compared to the control sample (Fig. 8f), also in keeping with the observation that *NRXN3* expression is reduced in PTHS neurons from patients #1 and #2 in qPCR and RNA-Seq experiments (Fig. 7b,c). Interestingly, in the PTHS samples, the decrease in intensity for the ∼40 kDa band (open arrowhead in Fig. 8f) is more pronounced than the drop in intensity for the ∼50 kDa band (filled arrowhead), which concurs with our observations that the ratio between the expression levels for the SE-NRXN3-i and SE-NRXN3-e splicing events decreases in patients #1 and #2 PTHS samples in comparison with control #1 (qPCR and rMATS-turbo results in Fig. 7e).

In combination, our results point to the higher prevalence of a distinct Neurexin protein isoform due to abnormal splicing patterns, potentially dictated by the dysregulated expression of RBFOX and NOVA1 splicing regulators, in neurons and organoids derived from Pitt-Hopkins Syndrome patients.

## DISCUSSION

RNA splicing plays a major role in producing the incredible variety of transcripts and their encoded protein products found in human cells. Yet, how the dysregulation of splicing events lead to disease is a topic that has only recently emerged. Over the last two decades, several studies have sought to elucidate how splicing can go from an important regulatory phenomenon able to regulate human gene expression and generate diverse mature mRNAs to pathogenic altered processes that can contribute to disease onset and progression. Of particular interest are splicing events that lead to exon skipping (ES), intron retention (IR), alternative 5’ or 3’ splice site usage (A5SS or A3SS), or the use of mutually exclusive exon^7^. Many of these alternative splicing events lead to frameshifts and premature stop codons that result in protein truncation (protein truncating variants; PTV), whose role in neurodevelopmental disorders are becoming increasingly clear^2^.

Even though the several recognized types of autism are the result of a complex interplay between genetic factors and environmental influences, their pathophysiology is still mostly unknown, and the disease mechanisms remain to be determined in most cases. It is becoming apparent that, among the many molecular features of this complex set of disorders, dysregulation of splicing is a more common phenomenon than initially anticipated^2^. One theme that is starting to be a topic of intense research in the field of autism molecular neurobiology is the decipherment of how altered splicing events or aberrant splicing patterns contribute to the cellular characteristics of the neural tissue in autistic children.

In this study, we determined that several genes exhibit aberrant splicing patterns in neurons derived from patients with a monogenic form of autism known as Pitt-Hopkins Syndrome (Figs. 3 and 4). We also showed that these altered splicing events are concomitant with the dysregulated expression of NOVA1 and RBFOX1/2/3 splicing regulators (Figs. 1 and 2). These are well known RNA binding proteins (RBPs) that facilitate spliceosome assembly and ensure accurate recognition of splice sites to lead to appropriate intron excision^52,53^. Extensive RBP-RNA interaction experiments have unveiled the participation of NOVA and RBFOX family members in splicing regulation, determining that they are prominently expressed in neurons and crucial to neurogenesis, axon guidance, and synapse formation^2^. An example of a neuronal gene regulated by alternative splicing is *DLG4* (encoding PSD-95), which may generate transcripts via the inclusion of specific exons that result in the incorporation of a domain lacking palmitoylation sites, thereby reducing membrane localization and enhancing excitatory synaptic transmission^54^.

Importantly, altered splicing in neurons of autistic patients impacts genes involved in neurodevelopment and synaptic plasticity, including synapse-related genes known to be risk factors for autism, such as *NLGN3*, *NLGN4*, *NRXN1*, *SHANK3*, and *CADPS*^55^. Our data showed that one of the aberrant splicing patterns observed in PTHS affect the *NRXN2* and *NRXN3* genes (Fig. 6), which encode Neurexin protein family members. It is interesting to note that altered splicing has been shown to affect the *NRXN1* gene in other forms of autism: these studies have pinpointed the existence of region-specific splicing of *NRXN1* in the prefrontal cortex, potentially influencing synapse formation^56^. Although this splicing event also involves a gene in the Neurexin family, our data show a different case in which a monogenic form of autism, caused by haploinsufficient heterozygous mutations in the *TCF4* gene, lead to changes in RBFOX and NOVA splicing regulators, potentially in turn leading to aberrant splicing patterns involving exon skipping events (Figs. 6 and 7), thereby producing a change in Neurexin 3 transcripts that modifies the balance between transmembrane and secreted forms of this synapse-related protein (Figs. 7 and 8).

Changes in NOVA and RBFOX expression have been studied by many groups over the last years, showing that these events disrupt neuronal development, leading to neurodevelopmental and neurodegenerative diseases^57^. Other studies further sought to understand how such dysregulation leads to pathological changes in splicing patterns. For example, RBFOX expression was shown to be downregulated in autism, a characteristic that leads to aberrant splicing in several ASD-associated genes in the SFARI database, including *SHANK3*, *CACNA1C*, and *TSC2^1^* and synapse-related genes *NRCAM* and *GRIN1*^58,59^. Moreover, mutations in *NOVA2* have been associated with autism and other neurodevelopmental disorders, impairing neurite outgrowth and axonal pathfinding^60,61^. Thousands of NOVA2 binding sites have been located in the mouse genome, particularly in genes involved in synapse and axon guidance such as *Dcc*, *Robo2*, *Slit2*, and *Epha5*^62,63^. Our study significantly expands these studies by describing that NOVA1 and RBFOX1/2/3 expression are impaired in a monogenic form of autism, PTHS, and by detecting thousands of genes in which splicing patterns are altered (Figs. 3 and 4).

It is noteworthy that many genes characterized by aberrant splicing patterns in PTHS are involved in synapse formation and organization (Fig. 4), which prompted us to focus on one particular splicing event (exon skipping) that affects the synapse-related *NRNX3* gene (Fig. 6). This finding may explain the impaired electrical activity we observed in the PTHS neural tissue^34^, as judged by the analysis of neurons in traditional 2D culture and in patient-derived brain organoids. These findings concur with our detection of transcript variants derived from alternative splicing in genes in the Neurexin family, which are known to control synapse formation during normal brain development^36,47^. In particular, the *NRXN3* gene encodes two major types of Neurexin, a large isoform encoded by longest isoform including most exons (α Neurexin-3 type) and a short isoform derived from transcription from an internal promoter (β type), both transmembrane-bound proteins^47^. Neurexins interact with Neuroligin proteins to control the rate at which synapses are formed in maturing neurons^64,65^.

Our study detected a change in the abundances of two transcripts derived from alternative splicing: in neurons of normal genotype, there seems to be an alternative splicing transcript that results from exon exclusion, leading to a α-type Neurexin lacking the transmembrane domain which is expected to be secreted^66^; in contrast, in PTHS neurons, the abundance of this secreted form is diminished relative to the transmembrane form (Fig. 8). Although the transcript with exon exclusion (SE-NRXN3-e) has already been described in the literature^66,67^, its involvement in the context of a neurodevelopmental disorder is novel and may open new avenues of investigation to understand how such splicing dysregulation leads to aberrant synapses and altered electrical activity in the neural tissue of autistic children. In the context of a neurodegenerative disorder (Alzheimer’s disease), the ratio between the Neurexin-3 transmembrane and soluble isoforms was found to be reduced^67^, which is exactly in opposition to what we observed in the context of autism. Further studies will illuminate the functional contribution of these ratio alterations in both diseases.

Another interesting possibility is that the changes in ratio between the transmembrane and secreted isoforms of Neurexin-3 may lead to differences in the content of GABAergic and glutamatergic synapses in PTHS. We raised this possibility based on the observation that deletion of the *Nrxn3* gene in mice led to a decrease in GABA-mediated inhibitory responses^65^. Outstandingly, we observed an augmentation of GABAergic neurons in PTHS cortical organoids, while these cells are mostly lacking in control cortical organoids (Fig. 2b,c,e,g,i), a piece of observation that may explain why PTHS cortical organoids have lower electrical activity^34^.

In conclusion, our study described alterations in the expression of splicing regulators in PTHS and how this is accompanied by aberrant splicing patterns affecting hundreds of genes involved in neurodevelopment. Further studies are needed to unravel how these alterations contribute to the pathology in autistic brains and to result in the identification of molecular therapeutic targets, eventually leading to the development of strategies to correct abnormalities in splicing regulation in autism.

## METHODS

### Patient-derived neural progenitor cell lines and neuronal differentiation

Our studies are based on RNA-Seq and single-cell RNA-Seq from our previous publication using neurons and brain organoids derived from patients with PTHS^34^. This study included subjects from volunteering families recruited through the University of Campinas (Supplementary Table 6) harboring heterozygous mutations in *TCF4* mutation. Details on the patients’ PTHS clinical symptoms are reported in Supplementary Table 6. Control subjects were the patients’ same-sex parents, with no prior history of psychiatric or genetic disorders. The participation of all subjects was approved by the Human Subjects Ethics Committees of the University of Campinas, under CAAE number 86136925.6.0000.5404. Written informed consent was obtained from all participating families after receiving a thorough description of the study and no compensation was provided to participants.

In our previous study, neural progenitor cells were obtained from reprogrammed iPSCs derived from skin fibroblasts subjected to cellular reprogramming via over-expression of the Yamanaka factors *OCT4*, *SOX2*, *KLF4*, and *MYC* (Cytotune iPS 2.0 Sendai reprogramming kit; Thermo Fisher Scientific)^34^. All iPSC lines were validated via immunostaining and SNP mapping to rule out the presence of unwanted chromosomal abnormalities and mutations. Results reported in this paper are from experiments conducted with one P15 iPSC clone per subject. Cultures were tested every two weeks for mycoplasma.

iPSC colonies maintained in mTeSR1 Plus medium (Stem Cell Technologies), followed by addition of 10 mM SB431542 and 1 mM dorsomorphin to form embryoid bodies. After 2 weeks of culturing, the embryoid bodies were plated onto Matrigel-coated plates with DMEM/F12 medium plus N2 and SM1 supplements (Stem Cell Technologies) and 20 ng/mL FGF-2. Neural rosettes were then picked and replated onto Matrigel-coated dishes, and the resulting NPCs were dissociated with Accutase and re-seeded onto plates coated with 10 μg/mL poly-ornithine (Sigma-Aldrich) and 5 μg/mL laminin (Thermo Fisher Scientific).

For neuronal differentiation, NPCs were seeded onto plates coated with poly-ornithine and laminin. At 90% confluency, medium was replaced with DMEM/F12 containing N2 and B27 supplements (Thermo Fisher Scientific) and 1% penicillin/streptomycin, with media changes occurring every 3 to 4 days.

### RNA sequencing from NPCs and neuronal cultures and single-cell RNA sequencing data from organoids

Our study employed bulk RNA sequencing data from NPCs and neurons derived from PTHS syndrome, which have been previously published by our group^34^. Briefly, these libraries were produced from NPCs of 4 subjects and 4 respective parental controls at passage 15 and from differentiating neuronal cultures of 3 parent-patient pairs and an additional patient line at 2 months in vitro. RNA was extracted from 3 independently prepared biological replicates, followed by Illumina library preparation using the stranded TruSeq kit (Illumina) and sequencing on an Illumina NovaSeq 6000 S4 instrument with 150 bp paired-end reads. Transcript-level expression abundance was estimated using Salmon (version 0.14.1) software^68^, with selective mapping (‘--validateMappings’) and correction for sequence-specific biases (‘--seqBias’), GC-content biases (‘--seqBias’), and fragment position bias (‘--posBias’). Reference transcripts for read mapping were obtained from the GENCODE 32 basic annotation^69^. MA plots of specific genes in RNA-Seq data was produced from dataframes containing the expression abundances in each individual library, using the ‘ggplot’ package for *R* language.

Single-cell RNA-Seq data used in this study was also previously published by our team (REF our paper), with samples from cortical organoids (CtOs) and subpallial organoids (sPOs).

Briefly, CtOs were produced from iPSC colonies dissociated using Accutase (Thermo Fisher Scientific), followed by neural induction in mTeSR1 Plus medium (Stem Cell Technologies) supplemented with 10 mM SB431542 (Stemgent) and 1 mM dorsomorphin (R&D Systems). Embryoid bodies formed were transferred to Neurobasal medium (Thermo Fisher Scientific) containing GlutaMAX, 1% B27 supplement (Thermo Fisher Scientific), 1% N2 NeuroPlex supplement (Gemini Bio-Products), 1% NEAA (Thermo Fisher Scientific), 1% penicillin/streptomycin (Thermo Fisher Scientific), 10 mM SB431542, and 1 mM dorsomorphin, for 7 days. NPC proliferation medium was then used, consisting of Neurobasal medium containing GlutaMAX, 1% B27, 1% NEAA, 20 ng/mL FGF-2 (Thermo Fisher Scientific) for 7 days, followed by 7 additional days with 20 ng/mL EGF (PeproTech). Neuronal differentiation was achieved in Neurobasal medium containing 1% GlutaMAX, 1% Gem21, 1% NEAA, 10 ng/mL of BDNF, 10 ng/mL of GDNF, 10 ng/mL of NT-3 (all from PeproTech), 200 mM L-ascorbic acid, and 1 mM dibutyryl-cAMP (Sigma-Aldrich), for 7 days. GABAergic-enriched subpallial organoids (sPOs) were producing using a similar protocol, with neural induction medium containing 5 μM IWP-2 (SelleckChem), from day 4 until day 10, followed by incubation in proliferation medium (Neurobasal medium containing 1% GlutaMAX, 1% Gem21, 1% NEAA, 20 ng/mL FGF-2, and 100 nM SAG (SelleckChem) for 7 days, followed by 2 additional days in the same medium supplemented with 20 ng/mL EGF (PeproTech).

Single cell RNA-Seq was performed on dissociated CtOs and GbOs at 8 weeks in vitro, with 15 organoids per library (5 organoids from each of three wells from independent experiments) dissociated with forceps and enzymatic digestion with Accutase. Libraries were prepared using the Chromium Single Cell 3’ v3 Library kit (10× Genomics) according to the manufacturer’s protocol, with a target of 20,000 cells per sample on the Chromium chip and PCR-amplification for a total of 10 cycles. Sequencing was performed on a NovaSeq 6000 S4 sequencer (Illumina) to produce 20,000 reads per single cell. Feature count matrices for each single-cell RNA-Seq library were generated separately using the ‘cellranger count’ command in the Cell Ranger software (version 4.0.0) and the GRCh38 2020-A reference dataset of human transcripts, yielding an estimated number of cells across all libraries in the range 2,944 to 6,466 per library, with a mean number of reads per cell ranging from 58,501 to 169,931. Cell type subpopulations were demarcated via automated annotation and refinement after manual inspection. Seurat (version 3.2.2)^70^ was used for producing UMAP and violin plots in this study, after normalization to 10,000 counts per cell and log-transformation with the ‘NormalizeData’ function, finding of variable features with the ‘FindVariableFunctions’ function, scaling with the ‘ScaleData’ function, and linear dimensional reduction (PCA) with the ‘RunPCA’ function. Non-linear reduction was done using the UMAP algorithm in the ‘RunUMAP’ function. For statistical comparisons of gene expression levels, we used Mann-Whitney U test.

### rMATS-turbo analysis of splicing events

rMATS-turbo analysis was conducted using control and patient RNA-seq data previously published by our group (GEO accession numbers GSE159392 and GSE189121)^34^. For alternative splicing analysis conducted with libraries with different read sequence lengths, sequences were trimmed down to the shortest length across all libraries (101 bp) using Trimmomatic software^71^. rMATS-turbo was run using the following arguments: end-pairing (‘-t’) equal to ‘paired’, read length (‘—readLength’) equal to 101 or 151, depending on the library, as well as a function to find novel splice-sites (‘—novelSS’). The raw output data was then filtered using rMATS-turbo specific in-built filtering script ‘rmats_filtering.py’^72^, with parameters set to filter events with a minimum average number of 10 read counts per sample, to filter out events with PSI < 0.05 or > 0.95, to filter events with FDR < 0.05, and to filter events with absolute values of IncLevelDifference (ΔPSI) > 0.05. Over-representation (ORA) and Gene-Set Enrichment Analysis (GSEA) were performed using the lists of genes for which SE events had absolute values of ΔPSI > 0.3, for each patient-control pair, using the WebGestalt software^73^ and Gene Ontology - Biological Process and KEGG pathway functional databases, with the ‘genome protein-coding’ reference set. For GSEA, the list of genes was ordered according to the absolute values of IncLevelDifference, with GO-Biological Process and KEGG pathway functional databases.

### RBP binding site recognition and enrichment analyses

For the analysis of motif enrichment for RBFOX and NOVA regulators, we recovered the genomic coordinates for the list of SE events that had absolute values of ΔPSI > 0.1, for patient-control pair #2. To recover the sequences of the exons involved in these SE events plus 200 bp of the neighboring introns, we subtracted 199 from the value of the initial exonic genomic coordinate (‘exonStart_0base’ column in the rMATS-turbo output table) and added 200 to the value of the final exonic genomic coordinate (‘exonEnd’ column). Sequences were recovered in fasta format using the ‘faidx’ function in samtools^74^ from the GRCh38 human genome database. RBP motif enrichment was conducted using the SEA software in the MEME suite^43^, with RNA motif database ‘Ray2013_rbp_Homo_sapiens.dna_encoded.meme’, which was modified to include motifs for NOVA1, RBFOX2, and RBFOX3. A position-specific scoring matrix (PSSM) for NOVA1 was obtained from the RBPmap developing team^75,76^; PSSM for RBFOX2 and RBFOX3 were produced using the ‘matrix2meme’ script in the MEME suite using published motif sequences from RNA Bind-n-Seq (RBNS) results (Supplementary Table S3 in ref^44^). SEA was run using default parameters plus arguments to avoid trimming (‘--notrim’) and the ‘--hofract’ parameter set to 0.5. As a negative control sequence set, we started with a random selection of 5,000 exons collected across all genes in the GRCh38 database using ‘awk’, ‘sort’, and ‘head’, to produce a FASTA file with exonic and neighboring intronic (200 bp) sequences.

As positive control sequences for RBFOX1/2/3 and NOVA1, we selected genes whose splicing is known to regulated by these proteins^2,77^ and recovered all exonic plus neighboring intronic (200 bp) sequences.

For the detection of binding sites of NOVA1 and RBFOX1/2/3 in specific genes (including *NRXN3*), we first obtained the FASTA files for the target genes from Ensembl (genome assembly GRCh38.p14; GCA_000001405.29) and then run the motif search using Find Individual Motif Occurrences (FIMO) software in the MEME suite^45^, with the ‘--norc’ argument to prevent motif scanning in the reverse complement strand. We ran this analysis for 50 genes at the top of the SE event list from rMATS-turbo, ordered according to the absolute values of ΔPSI. In parallel, we ran FIMO scanning for 50 random genes that were not found in the rMATS-turbo output SE event list. The frequencies of RBP binding sites were then normalized to 100,000 bp, followed by enrichment analysis using unpaired *t-*test in GraphPad Prism.

### PCR and sequencing of cloned bands

PCR was conducted using GoTaq DNA polymerase (Promega; cat# M3001) with primers p6 (5’ TCTGCTGAATGTTCAAGTG 3’) and p7 (5’ TCTTCAGTTTGGTGCTGTC 3’) for the first reaction and primers p6 and p8 (5’ GGAGATGGATGACAGCAC 3’) for the second reaction (Fig. 7f,g), using the following cycling parameters: 94°C 5 min, and 38 cycles of 94°C 45 s, 58°C 45 s, and 72°C 2 min. The fragments were then cloned into the pGEM-T-easy vector (Promega) and Sanger sequenced using universal M13 primer. Alignment was performed using Benchling alignment tool.

### Protein secondary structure analysis

Protein secondary structure analysis was performed using InterProScan^49^ for general domain prediction using the coding amino acid sequence of *NRXN3* ENST00000335750.7 and ENST00000554738.5 transcripts, representing the transmembrane and soluble isoforms, respectively, using default parameters. Specific transmembrane domain prediction was performed using DeepTMHMM 1.0 software^50^, inputting the same sequences mentioned above. The likelihood of secretion was evaluated using the SecretomeP (v 2.0) tool

### Real-time quantitative qPCR

For real-time qPCR, neuronal cultures were differentiated from NPCs seeded onto 6-well plates, followed by withdrawal of FGF-2 for 3 weeks, prior to RNA extraction with the RNeasy Mini Plus kit (Qiagen). All samples were analyzed with 3 independent biological replicates. cDNA was synthesized from total RNA using the Improm II First-Strand Reverse Transcription System (Promega). Real-time quantitative PCR (RT-qPCR) was performed using regular primers for SYBR Green technology, on an Applied Biosystems Real Time PCR system, with the following cycling parameters: 94°C for 3 min, followed by 40 cycles of 94°C for 30 s and 68°C for 1 min.

Amplification and denaturation curves for all probes were analyzed to verify amplification of just one amplicon. Normalization was achieved with endogenous control gene *ACTB*, and relative expression was calculated using the traditional ΔΔCt method. The following qPCR primers were used to detect all *NRXN3* transcript variants: p1 (5’ ACAACCCGTAAGAATGCGTC 3’) and p2 (5’ CATCACTTGAACATTCAGCAG 3’). The following variant-specific qPCR primers were used: p1, p3 (5’ GGAAAGACTCTTATCTGTACTC 3’), p4 (5’ TAGAGCTTCTGGCTGTACTC 3’), and p5 (5’ CGTAAGAATGCGCTCTACAG 3’).

### Cortical organoid derivation

For the generation of cortical brain organoids, we used our previously published protocol. iPSC colonies were dissociated using Accutase (Thermo Fisher Scientific; diluted with an equal volume of 1× PBS) for 12 min at 37 °C, followed by centrifugation for 3 min at 150 × *g*. Next, dissociated cells were resuspended in mTeSR Plus medium (Stem Cell Technologies) with 10 mM SB431542 (Stemgent) and 1 mM dorsomorphin (R&D Systems), followed by seeding 3–4 million cells onto low-binding 6-well plates. Organoids were cultured for 1 month on a shaker inside a CO_2_ incubator at 95 rpm. Rho kinase inhibitor (Y-27632; Calbiochem, Sigma-Aldrich) was added at a 5 mM concentration during the first 24 hours, in mTeSR Plus medium, which was supplemented with 10 mM SB431542 and 1 mM dorsomorphin; the same medium without Rho kinase inhibitor continued to be used for an additional 2 days. Next, medium was replaced with Neurobasal medium (Thermo Fisher Scientific) containing GlutaMAX, B27 supplement (Thermo Fisher Scientific), 1% N2 supplement (Thermo Fisher Scientific), 1% NEAA (Thermo Fisher Scientific), 1% penicillin/streptomycin (Thermo Fisher Scientific), 10 mM SB431542, and 1 mM dorsomorphin, for 5 days, with the addition of Rho kinase inhibitor on the first day of neural induction phase. Medium was then replaced with Neurobasal medium containing GlutaMAX, B27, 1% NEAA, penicillin/streptomycin, and 20 ng/mL FGF-2 (Thermo Fisher Scientific) for 8 days, followed by 6 additional days in the same medium further supplemented with 20 ng/mL EGF (PeproTech). Next, we used a transition phase by incubating the organoids in Neurobasal medium containing 1% GlutaMAX, 1% B27, 1% NEAA, penicillin/streptomycin for 3 days, followed by the neuronal differentiation and organoid maturation phase in Neurobasal medium containing 1% GlutaMAX, 1% B27, 1% NEAA, penicillin/streptomycin, 10 ng/mL of BDNF, 10 ng/mL of GDNF, and 10 ng/mL of NT-3 (all from PeproTech), for the remainder of the derivation protocol.

### Western Blotting

For Western Blotting analysis of Neurexin-3 isoforms, neuronal cultures were differentiated from NPCs seeded onto 6-well plates, followed by withdrawal of FGF-2 for 3 weeks, prior to protein extraction in RIPA buffer (0.05 M Tris-Cl pH 7.4; 1 M NaCl; 1% SDS) containing Protease Inhibitor Cocktail (Thermo Fisher Scientific), with mechanical dislodging of cells with a cell scraper. We also used organoid samples at 1 month in vitro, which were crushed in RIPA buffer containing protease inhibitor cocktail with the help of a plastic pistil. The crude extracts were clarified by centrifugation at 10,000 × *g* for 5 min, followed by discontinuous SDS-PAGE separation of proteins using a 7% acrylamide resolving gel, with 30 μg of total protein per sample. After protein transfer onto nitrocellulose membranes (Hybond-C; Amersham) using a TransBlot SD semi-dry transfer cell (BioRad), the membrane was blocked using TBS buffer (20 mM Tris-Cl pH 7.4; 150 mM NaCl) containing 0.1% Tween-20 (Sigma) and 3% BSA (Sigma), followed by detection of Neurexin-3 target proteins by incubation with primary antibody for 16 h at 4°C (anti-Neurexin-3 (E7K3G) rabbit monoclonal antibody; #48004S; Cell Signaling Technology; 1:1000) and fluorescent secondary antibody (IRDye 800CW goat anti-rabbit IgG. LiCOR; 1:5,000). Development was performed on an iBright CL1500 Imaging System (Thermo Fisher Scientific). As endogenous control, we used a mouse anti-GAPDH primary antibody (#60004; Proteintech; 1:2000) and secondary detection with IRDye 680CW goat anti-mouse antibody (LiCOR; 1:5,000).

### Statistical analysis and reproducibility

Data are presented as mean + SEM, unless otherwise indicated. Different types of statistical test were used throughout the study, including comparisons of means between two groups (PTHS against parent) in experiments that measured relative expression levels, or expression abundances used two-sample Welch’s t test, assuming unequal variances and heteroskedasticity. For comparing mean gene expression in single cell RNA-Seq data between two samples, we used the non-parametric Mann–Whitney U-test. Sample sizes are indicated in the figure legends. *P-*values are reported as asterisks in the figures for significance levels defined as *p*<0.05 (*), *p*<0.01 (**), or *p*<0.001 (***). Supplementary Table S8 presents extended results for all statistical tests performed, including sample sizes, statistical tests employed, effect sizes, statistics metrics (*H, F, t*, or *W*), along with exact *p*-values, listed according to the order of appearance in figure panels throughout the study. The numbers of replicates are also indicated in the figure legends and all attempts at replication were successful. When experiments involved data collected from independent biological replicates, results were collected from randomly chosen replicates (wells/plates of experiments involving cells in 2D culture) and blinding was used for most analyses comparing patients and control samples, including immunostaining. Blinding was not used when analyzing results from RNA sequencing and single cell RNA sequencing experiments, due to the inherently unbiased nature of the bioinformatic approaches used for quantitating gene expression and determining differential expression between genotypes or cell types. Statistical analyses were performed using Prism software (GraphPad; version 9.2.0). Figures were composed with Prism and Illustrator CS4 (Adobe; version 14.0.0).

## Materials Availability

Further information and requests for resources and reagents should be directed to and will be fulfilled by the corresponding author, Fabio Papes, upon reasonable request. When provided to others, unique reagents generated in this study will be available with a completed Materials Transfer Agreement.

## Data Availability

Data supporting the findings in this study are included within the Supplementary Material. RNA sequencing and single cell RNA sequencing raw and processed data used in this study are deposited at the Gene Expression Omnibus (GEO) of the National Center for Biotechnology Information (NCBI), under accession numbers GSE159392 (https://www.ncbi.nlm.nih.gov/geo/query/acc.cgi?acc=GSE159392), GSE159859 (https://www.ncbi.nlm.nih.gov/geo/query/acc.cgi?acc=GSE159859), GSE159860 (https://www.ncbi.nlm.nih.gov/geo/query/acc.cgi?acc=GSE159860), and GSE189121 (https://www.ncbi.nlm.nih.gov/geo/query/acc.cgi?acc=GSE189121), which are available publicly without restriction.

## Code Availability

No new custom software or code has been used in this paper. Codes (R programming language) used for bioinformatic analyses are all strictly based on pre-existing, regularly used published codes for RNA-Seq and single-cell RNA-Seq analyses, as described above.

## Acknowledgments

We thank Drs. Gonçalo Pereira, J. Andrés Yunes, and Maria Andreia Delbin for resources, Ana Paula Ferreira and Welbe Bragança for administrative and technical help, Dr. Elsa Molina (former director of the UCSD Stem Cell Genomics Core) for technical assistance with single cell RNA-Seq (made possible in part by CIRM Major Facilities grant #FA1-00607 to the Sanford Consortium for Regenerative Medicine). This work was supported by Sao Paulo Research Foundation (FAPESP) grants (#2020/11451-7 and #2022/12762-1), by FAPESP fellowships (#2024/18734-5 to P.K.K.F., #2024/12825-9 to B.B.H., #2025/09026-0 to J.G.B., #2024/20077-2 to G.D.V., and #2020/16525-9 to I.C.V.), and by a CNPq fellowship (L.L.D.). We would also like to thank Yael Mandel-Gutfreund and Dina Alexandrovich (RBPmap team^76^) for providing the NOVA1 motif PSSM.

## Author contributions

Conceptualization, F.P.; Methodology, F.P., P.K.K.F., A.P.C., and M.C.; Investigation, P.K.K.F., L.L.D., B.B.H., J.G.B., G.D.V., I.C.V., and M.H.B.; Writing – Original Draft, F.P., P.K.K.F.; Writing – Review & Editing, F.P. and P.K.K.F.; Funding Acquisition, F.P.; Resources, F.P., M.H.B., and M.C.; Supervision, F.P.

---

**Supplementary Figure S1:**
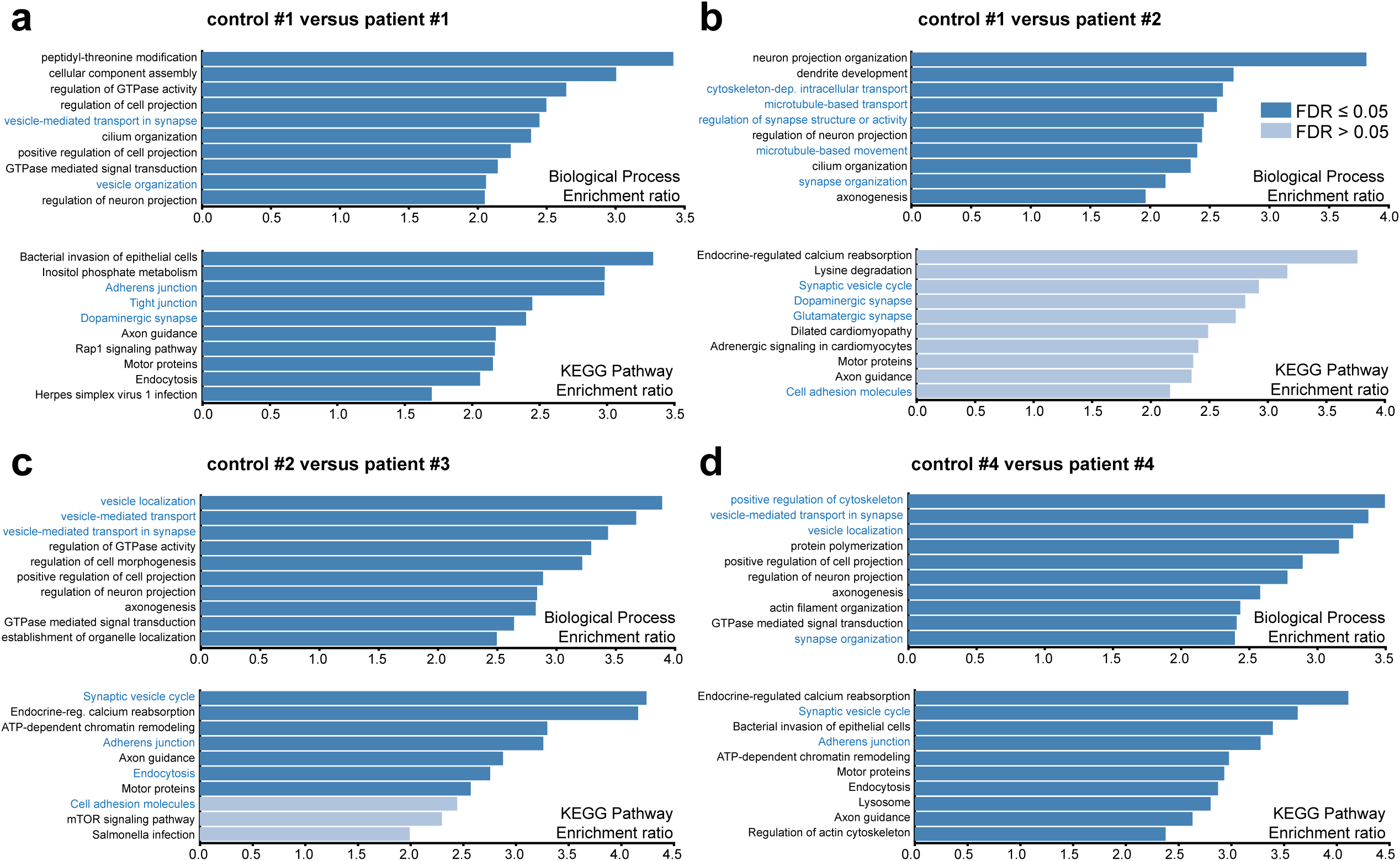
Over-representation analysis of genes involved in splicing events differentially used in PTHS. Enrichment ratios observed in Over-Representation Analysis (ORA) of splicing events with an absolute value of DPSI larger than 0.3 in the patient-control #2 pair, subjected to either Gene Ontology – Biological Process (top in each panel) or KEGG Pathway analysis (bottom in each panel). Only the 10 categories with the largest enrichment ratios are shown. The color of each bar represents the false discovery rate (FDR) and the category names indicated in blue represent those linked to synapse formation and function or cell-cell communication. The patient-control pairs shown in this figure complement the data for pair #2 presented in Fig. 4d,e.

**Supplementary Figure S2:**
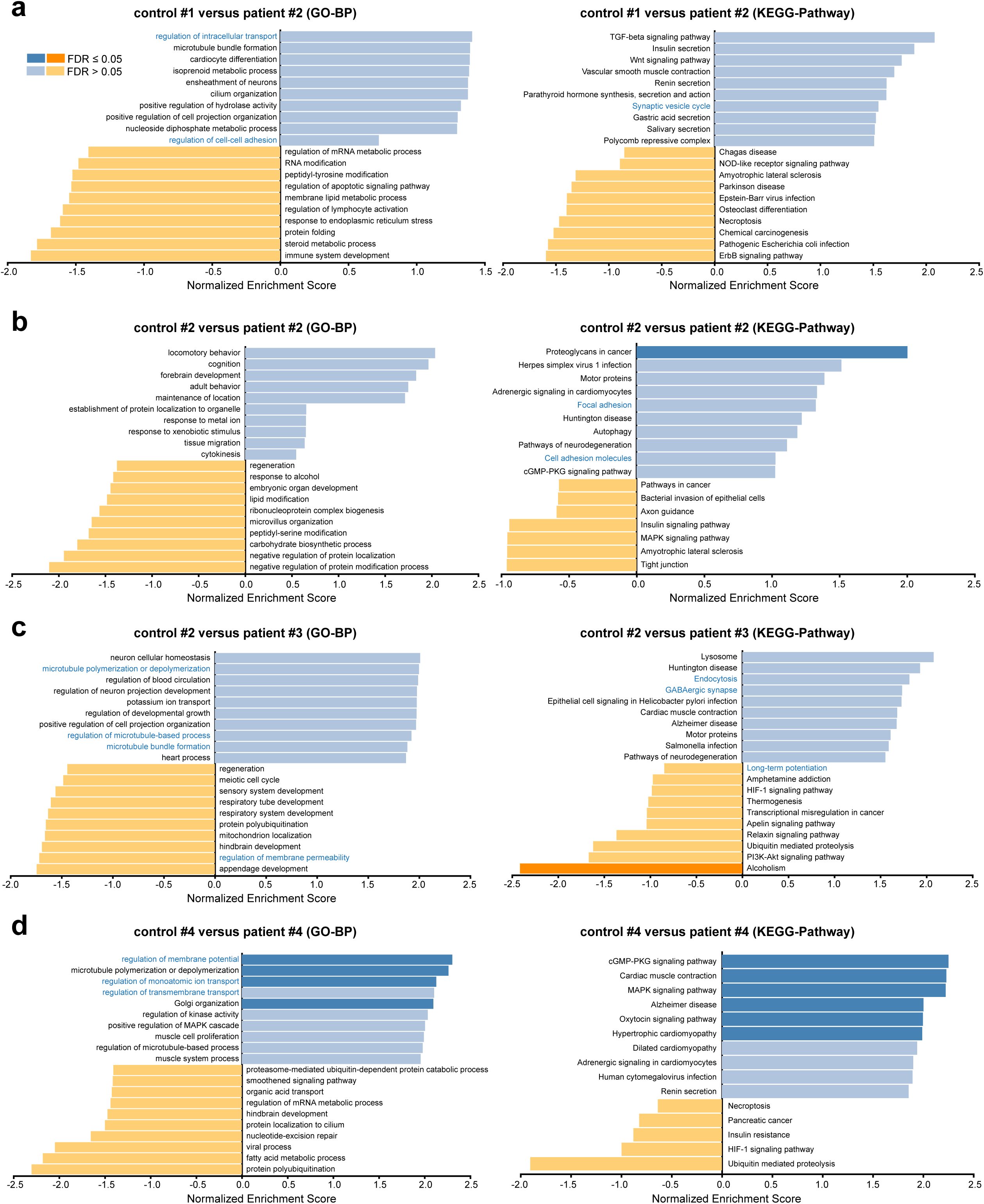
Gene-set enrichment analysis of genes involved in splicing events differentially used in PTHS. Enrichment ratios observed in Gene-Set Enrichment Analysis (GSEA) of splicing events with an absolute value of DPSI larger than 0.3 in the patient-control #2 pair, subjected to either Gene Ontology – Biological Process (left in each group) or KEGG Pathway analysis (right in each group). Only the 10 categories with the largest enrichment ratios are shown. The color of each bar represents the false discovery rate (FDR) and the category names indicated in blue represent those linked to synapse formation and function or cell-cell communication. The patient-control pairs shown in this figure complement the data for pair #2 presented in Fig. 4f,g.

**Supplementary Figure S3:**
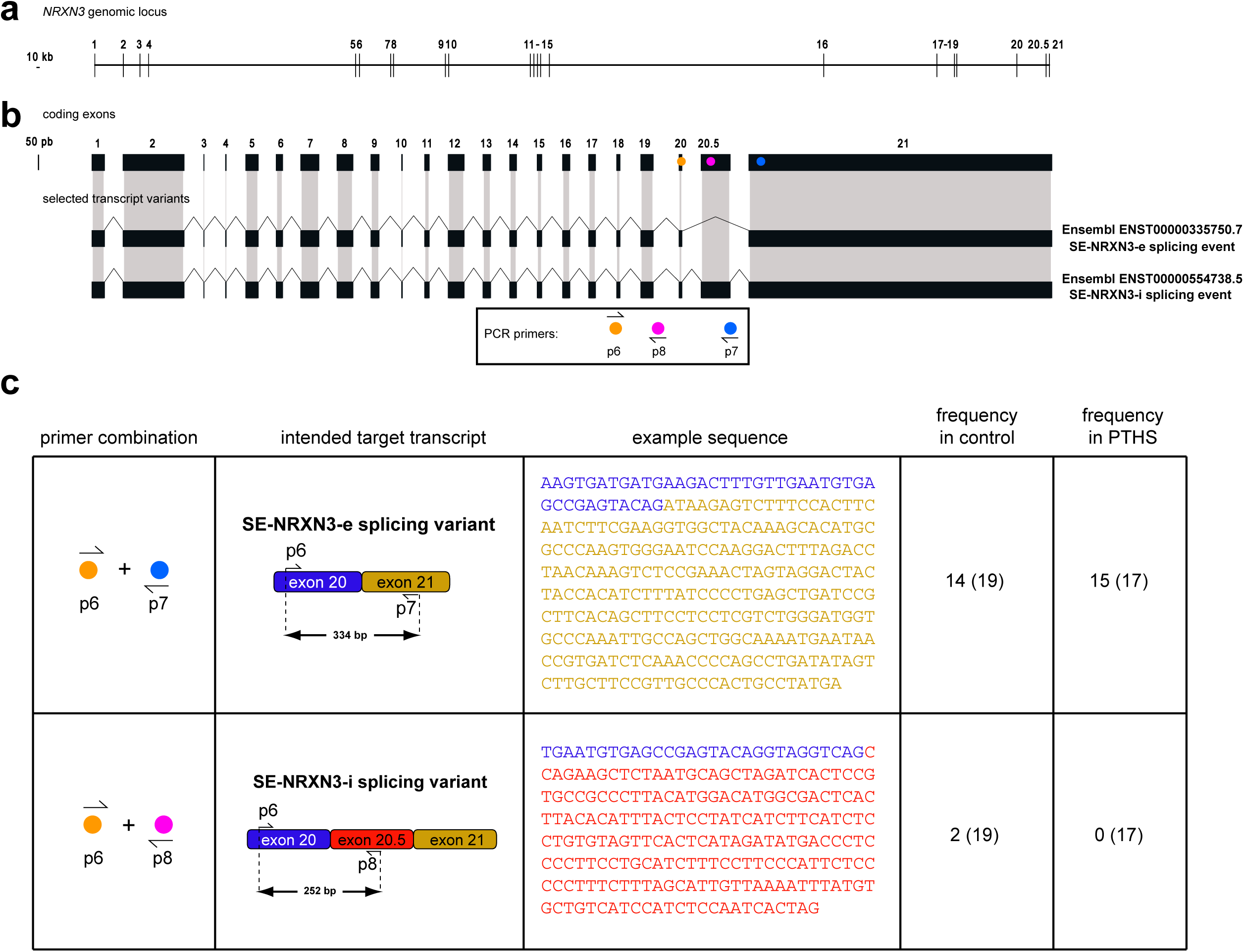
Supporting data for the analysis of splicing patterns in the *NRXN3* gene via PCR and Sanger sequencing. **(a)** Structure of the *NRNX3* (Neurexin-3) genomic locus, with the distribution of exons. Notice that exon 20.5 is annotated as an additional exon between 20 and 21, because it is not part of the transcript exhibited as the canonical *NRXN3* transcript in Ensembl. **(b)** Schematic representation of the coding exons in the *NRXN3* gene, with the location of PCR oligos to amplify the exon-exon junctions: primers p6 and p7 amplify the transcript variant without exon 20.5 (SE-NRXN3-e), while primers p6 and p8 amplify the transcript variant including exon 20.5 (SE-NRXN3-i). Exons are shown to scale. **(c)** Representation of the primer combinations and expected amplicon sizes, as well as an example of amplicon sequence. The frequencies of each amplicon in control and PTHS samples are shown in the last two columns (numbers in parentheses represent the total number of amplicons sequenced).

## Supplementary Material

### Supplementary Table Legends

**Supplementary Table S1: Expression abundances of selected genes in RNA-Seq data from PTHS and control neurons and neural progenitor cells.** The expression abundances in independent biological replicates (A, B, and C) for each subject, in transcripts per million reads (TPM), are shown for a selection of splicing regulators in the NOVA and RBFOX families. The means of expression in each subjects and log fold-changes (in base 2) for the comparison between each subject and a control are also shown.

**Supplementary Table S2: Splicing pattern analysis results using rMATS-turbo**. Each tab contains the filtered rMATS-turbo output for one patient versus control pair. Different tabs also indicate the results for the five types of alternative splicing: skipped exon (SE), alternative 5′ splice site (A5SS), alternative 3′ splice site (A3SS), mutually exclusive exon (MXE), and retained intron (RI). The common columns in all tabs are as follows: “ID”: rMATS event ID; “GeneID”: Ensembl gene ID; “geneSymbol”: gene name; “chr”: chromosome; “strand”: strand of the gene; “IJC_SAMPLE_1”: number of counts that support the inclusion of the splicing event, for the control sample (replicates are comma-separated); “SJC_SAMPLE_1”: number of counts that support the exclusion of the splicing event, for the control sample (replicates are comma-separated); “IJC_SAMPLE_2”: number of counts that support the inclusion of the splicing event, for the patien sample (replicates are comma-separated); “SJC_SAMPLE_2”: number of counts that support the exclusion of the splicing event, for the patient sample (replicates are comma-separated); “IncFormLen”: length of inclusion form, used for normalization purposes; “SkipFormLen”: length of skipping form, used for normalization purposes; “PValue”: significance of splicing difference between the two sample groups; “FDR”: false discovery rate; “IncLevel1”: inclusion level (in percentage) for the control sample (replicates are comma-separated), calculated from normalized counts; “IncLevel2”: inclusion level (in percentage) for the patient sample (replicates are comma-separated), calculated from normalized counts; “IncLevelDifference”: average(IncLevel1) − average(IncLevel2). See also the meaning of other columns, which indicate the genomic coordinates of the exons involved, in the rMATS-turbo original publication^1^.

**Supplementary Table S3: Common splicing events (and respective genes) detected by rMATS-turbo across all patient-control pairs.** Each of the first 5 tabs lists the common splicing events in each of the five categories of alternative splicing (SE, RI, MXE, A5SS, and A3SS). The meaning of each column is the same as in Supplementary Table S2. The last 5 tabs list the genes corresponding to these common splicing events.

**Supplementary Table S4: List of genes known to be regulated by NOVA1 and RBFOX1/2/3 splicing regulators**. Each tab lists the genes for which we extracted genomic sequences to produce the positive control datasets used for the Site Enrichment Analysis (SEA), for NOVA1 and RBFOX1/2/3. In the latter case, we combined information pertaining to genes known to be regulated by RBFOX1, RBFOX2, and/or RBFOX3 into a single list.

**##Supplementary Table S5: Lists of genes detected by rMATS-turbo in our PTHS splicing analysis and lists of genes binding NOVA and RBFOX splicing regulators**. The first tab lists the genes for which our rMATS-turbo analysis detected aberrant splicing events, using a threshold of absolute values of ΔPSI above 0.1. The last three tabs contain lists of genes detected by HITS-CLiP in published datasets, focusing on NOVA1^2,3^ or RBFOX1/2/3^4^. These lists were used to produce the Venn diagrams in Fig 5f-h.

**Supplementary Table S6: Summary of participating subjects and clinical characteristics.** For all patients, controls are parents of matching sex. Abbreviations: FG: dysmorphic facial *gestalt*; SM/MM: severe (SM) or mild (MM) motor delay at age 3; AS: absent speech; CT: constipation; SZ: seizures; BA: breathing problems (hyperventilation or apnea); UA: urinary abnormalities (retention or incontinence); VA: visual abnormalities (ocular anomalies); RB: repetitive behaviors; abMRI: brain anomalies (thinned *corpus callosum*) detected by MRI.

**Supplementary Table S7: Motifs for NOVA1 and RBFOX1/2/3 used for motif search and enrichment analyses.** The RBP motifs for NOVA1, RBFOX1, RBFOX2, and RBFOX3 are shown in tabular PSSM (Position-Specific Scoring Matrix) format.

**Supplementary Table S8: Summary of statistical tests and metrics.** For all figure panels (listed in order of appearance in the paper), we present complete statistical evaluation results, including type of analysis, groups being compared, means and standard errors of the means (SEM) for each group, numbers of subjects, independent replicates, technical replicates, or numbers of cells, as well as the number of observations used to calculate means and perform statistical comparisons (N), the statistical test used, metrics results (t, H, W, or F), effect sizes, exact p-values for global statistical comparisons, and, when necessary, effect sizes and exact p-values for each individual pairwise comparison after post-hoc test. When experimentation involved more than one independent replicate per subject cell line, or more than one technical replicate per independent biological replicate, these numbers are also indicated in separate columns, even though the statistical test was computed based solely on the means of different lines in these cases. The comparisons of gene expression between control and PTHS single cell RNA-Seq libraries are listed in the second tab (part 2), including gene name, cell types, groups being compared, means and SEM, fold change (PTHS versus control), as well as statistical test results.

## References

1. Weyn-Vanhentenryck, S. M. et al. HITS-CLIP and Integrative Modeling Define the Rbfox Splicing-Regulatory Network Linked to Brain Development and Autism. Cell Rep 6, 1139–1152 (2014).

2. Jeong, J., Yoo, H. J., An, J. Y. & Jeong, S. Dysregulated RNA-binding proteins and alternative splicing: Emerging roles in autism spectrum disorder. Mol Cells 48, 100237 (2025).

3. Kjer-Hansen, P. & Weatheritt, R. J. The function of alternative splicing in the proteome: rewiring protein interactomes to put old functions into new contexts. Nature Structural & Molecular Biology 2023 30:12 30, 1844–1856 (2023).

4. Zheng, S. Alternative splicing programming of axon formation. Wiley Interdiscip Rev RNA 11, e1585 (2020).

5. Quesnel-Vallières, M., Weatheritt, R. J., Cordes, S. P. & Blencowe, B. J. Autism spectrum disorder: insights into convergent mechanisms from transcriptomics. Nature Reviews Genetics 2018 20:1 20, 51–63 (2018).

6. Kim, K. K., Adelstein, R. S. & Kawamoto, S. Identification of neuronal nuclei (NeuN) as Fox-3, a new member of the Fox-1 gene family of splicing factors. Journal of Biological Chemistry 284, 31052–31061 (2009).

7. Jacko, M. et al. Rbfox Splicing Factors Promote Neuronal Maturation and Axon Initial Segment Assembly. Neuron 97, 853–868.e6 (2018).

8. An, J. Y. et al. Towards a molecular characterization of autism spectrum disorders: An exome sequencing and systems approach. Transl Psychiatry 4, (2014).

9. Teplova, M. et al. Protein-RNA and protein-protein recognition by dual KH1/2 domains of the neuronal splicing factor Nova-1. Structure 19, 930–944 (2011).

10. Scala, M. et al. De novo truncating NOVA2 variants affect alternative splicing and lead to heterogeneous neurodevelopmental phenotypes. Hum Mutat 43, 1299–1313 (2022).

11. Licatalosi, D. D. et al. HITS-CLIP yields genome-wide insights into brain alternative RNA processing. Nature 456, 464–469 (2008).

12. O’Leary, A. et al. Behavioural and functional evidence revealing the role of RBFOX1 variation in multiple psychiatric disorders and traits. Mol Psychiatry 27, 4464 (2022).

13. Brockschmidt, A. et al. Severe mental retardation with breathing abnormalities (Pitt-Hopkins syndrome) is caused by haploinsufficiency of the neuronal bHLH transcription factor TCF4. Hum Mol Genet 16, 1488–94 (2007).

14. Amiel, J. et al. Mutations in TCF4, Encoding a Class I Basic Helix-Loop-Helix Transcription Factor, Are Responsible for Pitt-Hopkins Syndrome, a Severe Epileptic Encephalopathy Associated with Autonomic Dysfunction. The American Journal of Human Genetics 80, 988–993 (2007).

15. Zweier, C. et al. Haploinsufficiency of TCF4 Causes Syndromal Mental Retardation with Intermittent Hyperventilation (Pitt-Hopkins Syndrome). The American Journal of Human Genetics 80, 994–1001 (2007).

16. Pitt, D. & Hopkins, I. A syndrome of mental retardation, wide mouth and intermittent overbreathing. Aust Paediatr J 14, 182–184 (1978).

17. Kim, H., Berens, N. C., Ochandarena, N. E. & Philpot, B. D. Region and Cell Type Distribution of TCF4 in the Postnatal Mouse Brain. Front Neuroanat 14, (2020).

18. de Pontual, L. et al. Mutational, functional, and expression studies of the TCF4 gene in Pitt-Hopkins syndrome. Hum Mutat 30, 669–76 (2009).

19. Jung, M. et al. Analysis of the expression pattern of the schizophrenia-risk and intellectual disability gene TCF4 in the developing and adult brain suggests a role in development and plasticity of cortical and hippocampal neurons. Mol Autism 9, 1–15 (2018).

20. Sepp, M., Kannike, K., Eesmaa, A., Urb, M. & Timmusk, T. Functional Diversity of Human Basic Helix-Loop-Helix Transcription Factor TCF4 Isoforms Generated by Alternative 5′ Exon Usage and Splicing. PLoS One 6, e22138 (2011).

21. Chen, E. S. et al. Molecular convergence of neurodevelopmental disorders. Am J Hum Genet 95, 490–508 (2014).

22. Schmidt-Edelkraut, U., Daniel, G., Hoffmann, A. & Spengler, D. Zac1 Regulates Cell Cycle Arrest in Neuronal Progenitors via Tcf4. Mol Cell Biol 34, 1020–1030 (2014).

23. Hill, M. J. et al. Knockdown of the schizophrenia susceptibility gene TCF4 alters gene expression and proliferation of progenitor cells from the developing human neocortex. Journal of Psychiatry and Neuroscience 42, 181–188 (2017).

24. Forrest, M. P. et al. The Psychiatric Risk Gene Transcription Factor 4 (TCF4) Regulates Neurodevelopmental Pathways Associated With Schizophrenia, Autism, and Intellectual Disability. Schizophr Bull 44, 1100–1110 (2018).

25. Page, S. C. et al. The schizophrenia-and autism-associated gene, transcription factor 4 regulates the columnar distribution of layer 2/3 prefrontal pyramidal neurons in an activity-dependent manner. Mol Psychiatry 23, 304–315 (2018).

26. Fischer, B. et al. E-proteins orchestrate the progression of neural stem cell differentiation in the postnatal forebrain. Neural Dev 9, 23 (2014).

27. Marangi, G. & Zollino, M. Pitt-Hopkins Syndrome and Differential Diagnosis: A Molecular and Clinical Challenge. J Pediatr Genet 4, 168–76 (2015).

28. Zollino, M. et al. Diagnosis and management in Pitt-Hopkins syndrome: First international consensus statement. Clin Genet 95, 462–478 (2019).

29. Stefansson, H. et al. Common variants conferring risk of schizophrenia. Nature 460, 744–747 (2009).

30. Ripke, S. et al. Genome-wide association study identifies five new schizophrenia loci. Nat Genet 43, 969–976 (2011).

31. Smoller, J. W. et al. Identification of risk loci with shared effects on five major psychiatric disorders: a genome-wide analysis. The Lancet 381, 1371–1379 (2013).

32. Wray, N. R. et al. Genome-wide association analyses identify 44 risk variants and refine the genetic architecture of major depression. Nat Genet 50, 668–681 (2018).

33. Gelernter, J. et al. Genome-wide association study of post-traumatic stress disorder reexperiencing symptoms in >165,000 US veterans. Nat Neurosci 22, 1394–1401 (2019).

34. Papes, F. et al. Transcription Factor 4 loss-of-function is associated with deficits in progenitor proliferation and cortical neuron content. Nat Commun 13, (2022).

35. Teixeira, J. R., Szeto, R. A., Carvalho, V. M. A., Muotri, A. R. & Papes, F. Transcription factor 4 and its association with psychiatric disorders. Transl Psychiatry 11, (2021).

36. Gomez, A. M., Traunmüller, L. & Scheiffele, P. Neurexins: Molecular Codes for Shaping Neuronal Synapses. Nat Rev Neurosci 22, 137 (2021).

37. Trujillo, C. A. et al. Complex Oscillatory Waves Emerging from Cortical Organoids Model Early Human Brain Network Development. Cell Stem Cell 25, 558–569.e7 (2019).

38. Yoon, S.-J. et al. Reliability of human cortical organoid generation. Nat Methods 16, 75–78 (2019).

39. Gasiorowska, A. et al. The Biology and Pathobiology of Glutamatergic, Cholinergic, and Dopaminergic Signaling in the Aging Brain. Front Aging Neurosci 13, 654931 (2021).

40. Sun, L. et al. Transcriptomic insights into fate choice of pallial versus subpallial GABAergic neurons. Nature Communications 16, 1–20 (2025).

41. Wang, Y. et al. rMATS-turbo: an efficient and flexible computational tool for alternative splicing analysis of large-scale RNA-seq data. Nature Protocols 2024 19:4 19, 1083–1104 (2024).

42. Wang, J., Vasaikar, S., Shi, Z., Greer, M. & Zhang, B. WebGestalt 2017: a more comprehensive, powerful, flexible and interactive gene set enrichment analysis toolkit. Nucleic Acids Res 45, W130–W137 (2017).

43. Bailey, T. L. & Grant, C. E. SEA: Simple Enrichment Analysis of motifs. bioRxiv 2021.08.23.457422 (2021) doi:10.1101/2021.08.23.457422.

44. Dominguez, D. et al. Sequence, Structure, and Context Preferences of Human RNA Binding Proteins. Mol Cell 70, 854–867.e9 (2018).

45. Grant, C. E., Bailey, T. L. & Noble, W. S. FIMO: scanning for occurrences of a given motif. Bioinformatics 27, 1017–1018 (2011).

46. Yuan, Y. et al. Cell type-specific CLIP reveals that NOVA regulates cytoskeleton interactions in motoneurons. Genome Biol 19, 1–19 (2018).

47. Zhang, R., Jiang, H. X., Liu, Y. J. & He, G. Q. Structure, function, and pathology of Neurexin-3. Genes Dis 10, 1908–1919 (2023).

48. Hishimoto, A. et al. Neurexin 3 polymorphisms are associated with alcohol dependence and altered expression of specific isoforms. Hum Mol Genet 16, 2880–2891 (2007).

49. Blum, M. et al. InterPro: the protein sequence classification resource in 2025. Nucleic Acids Res 53, D444–D456 (2025).

50. Hallgren, J. et al. DeepTMHMM predicts alpha and beta transmembrane proteins using deep neural networks. bioRxiv 2022.04.08.487609 (2022) doi:10.1101/2022.04.08.487609.

51. Teufel, F. et al. SignalP 6.0 predicts all five types of signal peptides using protein language models. Nat Biotechnol 40, 1023–1025 (2022).

52. Kumari, A., Sedehizadeh, S., Brook, J. D., Kozlowski, P. & Wojciechowska, M. Differential fates of introns in gene expression due to global alternative splicing. Hum Genet 141, 31–47 (2022).

53. Jeong, S. SR Proteins: Binders, Regulators, and Connectors of RNA. Mol Cells 40, 1–9 (2017).

54. Han, K. & Kim, E. Synaptic adhesion molecules and PSD-95. Prog Neurobiol 84, 263–283 (2008).

55. Quesnel-Vallières, M., Weatheritt, R. J., Cordes, S. P. & Blencowe, B. J. Autism spectrum disorder: insights into convergent mechanisms from transcriptomics. Nat Rev Genet 20, 51–63 (2019).

56. Fuccillo, M. V. et al. Single-Cell mRNA Profiling Reveals Cell-Type-Specific Expression of Neurexin Isoforms. Neuron 87, 326–340 (2015).

57. Parra, A. S. & Johnston, C. A. Emerging Roles of RNA-Binding Proteins in Neurodevelopment. J Dev Biol 10, (2022).

58. An, J. Y. et al. Towards a molecular characterization of autism spectrum disorders: an exome sequencing and systems approach. Transl Psychiatry 4, (2014).

59. Voineagu, I. et al. Transcriptomic analysis of autistic brain reveals convergent molecular pathology. Nature 474, 380–386 (2011).

60. Mattioli, F. et al. De Novo Frameshift Variants in the Neuronal Splicing Factor NOVA2 Result in a Common C-Terminal Extension and Cause a Severe Form of Neurodevelopmental Disorder. Am J Hum Genet 106, 438–452 (2020).

61. Scala, M. et al. De novo truncating NOVA2 variants affect alternative splicing and lead to heterogeneous neurodevelopmental phenotypes. Hum Mutat 43, 1299–1313 (2022).

62. Licatalosi, D. D. et al. HITS-CLIP yields genome-wide insights into brain alternative RNA processing. Nature 456, 464–469 (2008).

63. Ule, J. et al. CLIP identifies Nova-regulated RNA networks in the brain. Science 302, 1212–1215 (2003).

64. Graf, E. R., Zhang, X., Jin, S. X., Linhoff, M. W. & Craig, A. M. Neurexins Induce Differentiation of GABA and Glutamate Postsynaptic Specializations via Neuroligins. Cell 119, 1013 (2004).

65. Südhof, T. C. Synaptic Neurexin Complexes: A Molecular Code for the Logic of Neural Circuits. Cell 171, 745 (2017).

66. Hishimoto, A. et al. Neurexin 3 polymorphisms are associated with alcohol dependence and altered expression of specific isoforms. Hum Mol Genet 16, 2880–2891 (2007).

67. Hishimoto, A. et al. Neurexin 3 transmembrane and soluble isoform expression and splicing haplotype are associated with neuron inflammasome and Alzheimer’s disease. Alzheimers Res Ther 11, (2019).

68. Patro, R., Duggal, G., Love, M. I., Irizarry, R. A. & Kingsford, C. Salmon provides fast and bias-aware quantification of transcript expression. Nat Methods 14, 417–419 (2017).

69. Frankish, A. et al. GENCODE reference annotation for the human and mouse genomes. Nucleic Acids Res 47, D766–D773 (2019).

70. Sun, A. X. et al. Potassium channel dysfunction in human neuronal models of Angelman syndrome. Science (1979) 366, 1486–1492 (2019).

71. Bolger, A. M., Lohse, M. & Usadel, B. Trimmomatic: a flexible trimmer for Illumina sequence data. Bioinformatics 30, 2114 (2014).

72. Wang, Y. et al. rMATS-turbo: an efficient and flexible computational tool for alternative splicing analysis of large-scale RNA-seq data. Nat Protoc 19, 1083–1104 (2024).

73. Elizarraras, J. M. et al. WebGestalt 2024: faster gene set analysis and new support for metabolomics and multi-omics. Nucleic Acids Res 52, W415–W421 (2024).

74. Danecek, P. et al. Twelve years of SAMtools and BCFtools. Gigascience 10, 1–4 (2021).

75. Paz, I., Argoetti, A., Cohen, N., Even, N. & Mandel-Gutfreund, Y. RBPmap: A Tool for Mapping and Predicting the Binding Sites of RNA-Binding Proteins Considering the Motif Environment. Methods Mol Biol 2404, 53–65 (2022).

76. Paz, I., Kosti, I., Ares, M., Cline, M. & Mandel-Gutfreund, Y. RBPmap: A web server for mapping binding sites of RNA-binding proteins. Nucleic Acids Res 42, (2014).

77. Piton, A. NOVA1/2 genes and alternative splicing in neurodevelopment. Curr Opin Genet Dev 93, (2025).

78. Abrahams, B. S. et al. SFARI Gene 2.0: a community-driven knowledgebase for the autism spectrum disorders (ASDs). Mol Autism 4, 36 (2013).

## Supplementary Material References

1. Wang, Y. et al. rMATS-turbo: an efficient and flexible computational tool for alternative splicing analysis of large-scale RNA-seq data. Nature Protocols 2024 19:4 19, 1083–1104 (2024).

2. Yuan, Y. et al. Cell type-specific CLIP reveals that NOVA regulates cytoskeleton interactions in motoneurons. Genome Biol 19, 1–19 (2018).

3. Ule, J. et al. CLIP identifies Nova-regulated RNA networks in the brain. Science 302, 1212–1215 (2003).

4. Weyn-Vanhentenryck, S. M. et al. HITS-CLIP and Integrative Modeling Define the Rbfox Splicing-Regulatory Network Linked to Brain Development and Autism. Cell Rep 6, 1139–1152 (2014).

