## Supplementary figures and images for "Abnormal expression of splicing regulators RBFOX and NOVA is associated with aberrant splicing patterns at the Neurexin-3 gene in a monogenic autism spectrum disorder"

### Supplementary Fig. 1

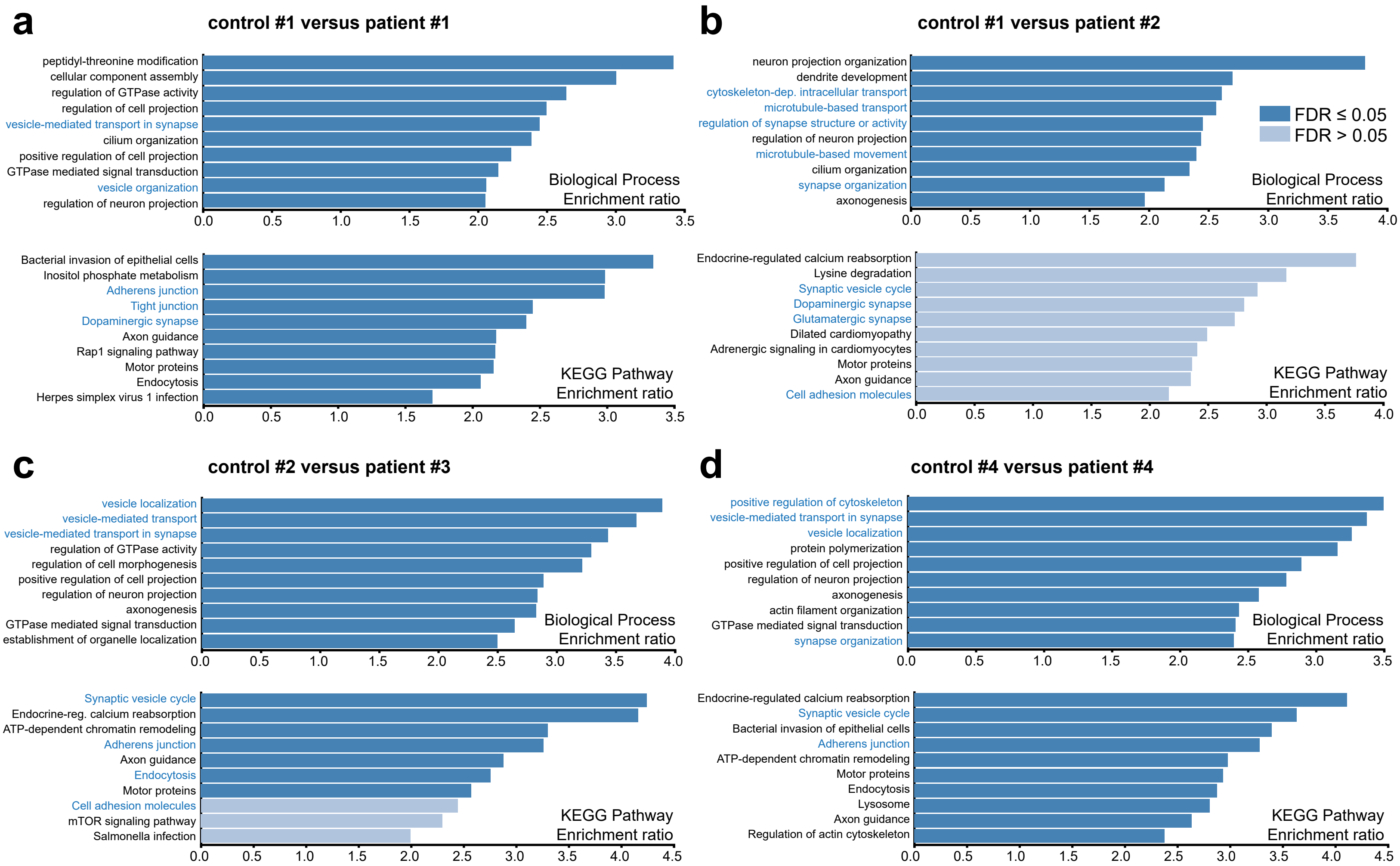

### Supplementary Fig. 2

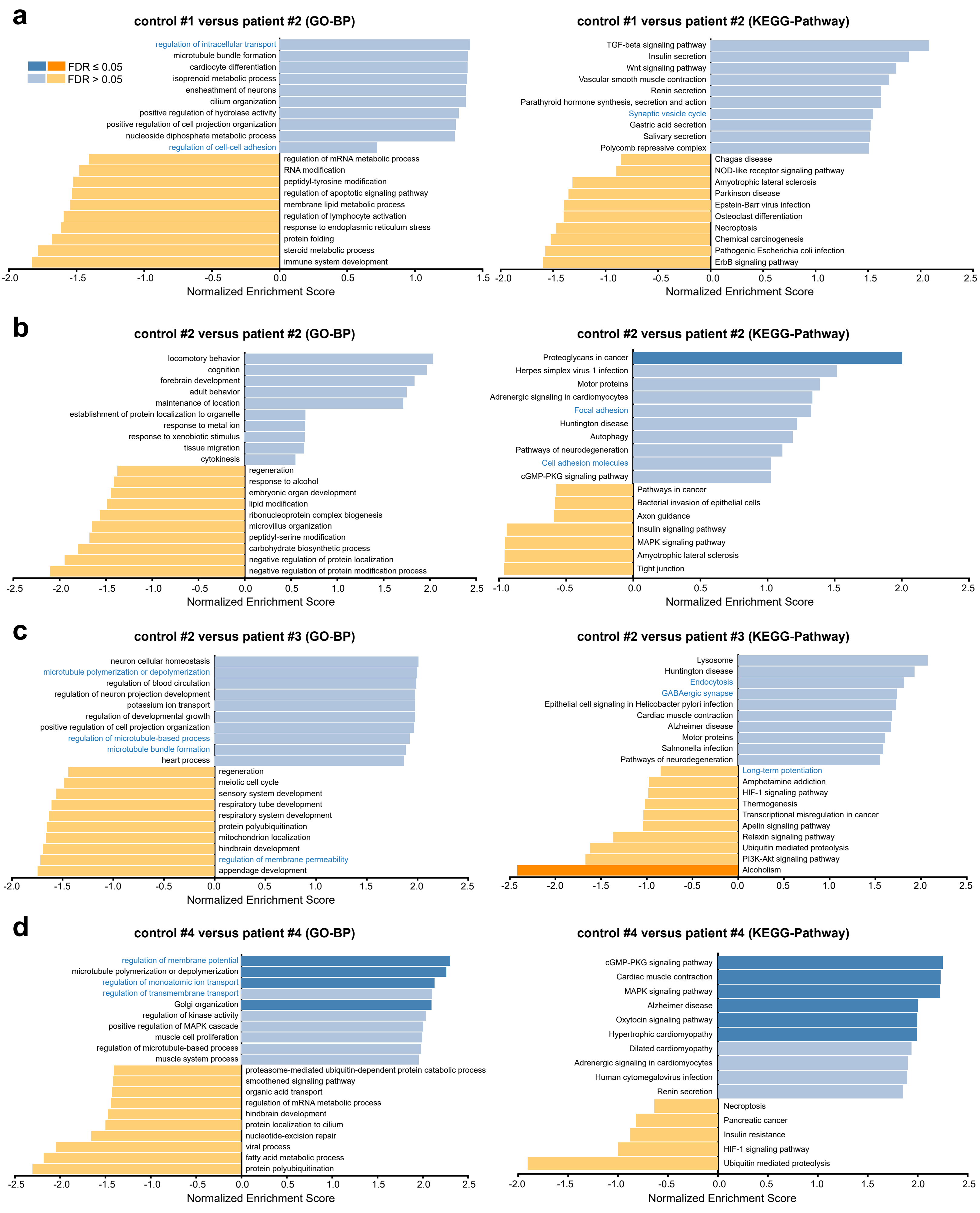
