## Supplementary Fig. 3 for "Abnormal expression of splicing regulators RBFOX and NOVA is associated with aberrant splicing patterns at the Neurexin-3 gene in a monogenic autism spectrum disorder"

### Figure S3

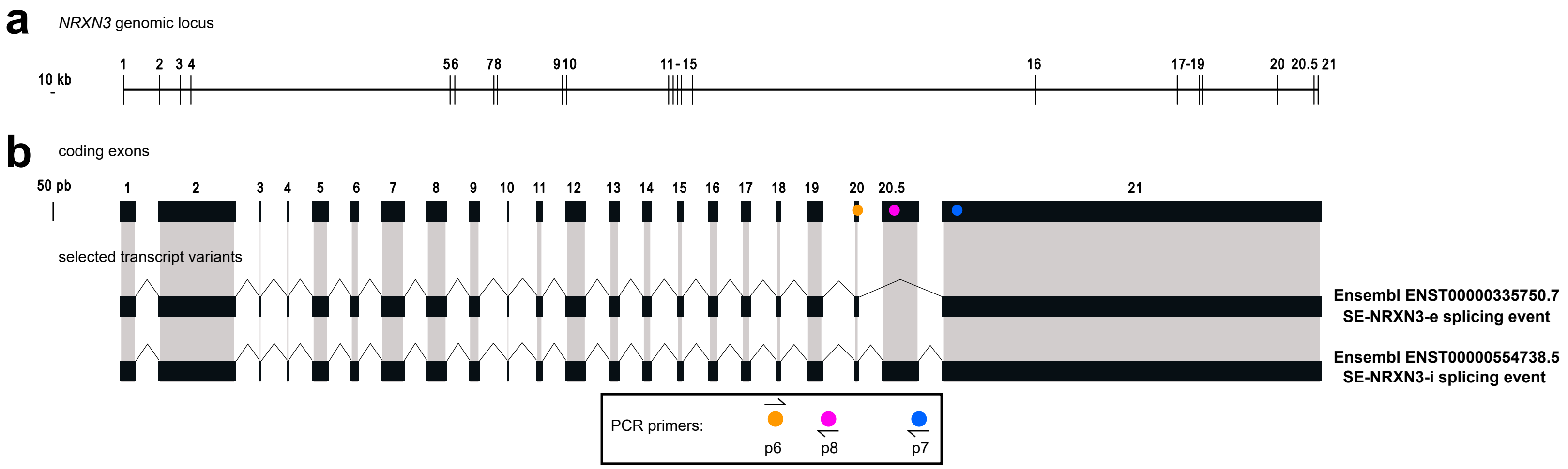


**c**

| primer combination | intended target transcript | example sequence | frequency in control | frequency in PTHS |
|---|---|---|---|---|
| 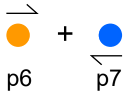 p6 + p7 | **SE-NRXN3-e splicing variant** 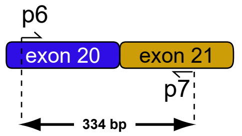 p6 exon 20 exon 21 p7 334 bp | AAGTGATGATGAAGACTTTGTTGAATGTGA<br>GCCGAGTACAGATAAGAGTCTTTCCACTTC<br>AATCTTCGAAGGTGGCTACAAAGCACATGC<br>GCCCAAGTGGGAATCCAAGGACTTTAGACC<br>TAACAAAGTCTCCGAAACTAGTAGGACTAC<br>TACCACATCTTTATCCCCTGAGCTGATCCG<br>CTTCACAGCTTCCTCCTCGTCTGGGATGGT<br>GCCCAAATTGCCAGCTGGCAAAATGAATAA<br>CCGTGATCTCAAACCCCAGCCTGATATAGT<br>CTTGCTTCCGTTGCCCACTGCCTATGA | 14 (19) | 15 (17) |
| 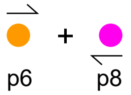 p6 + p8 | **SE-NRXN3-i splicing variant** 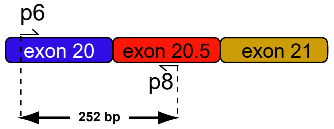 p6 exon 20 exon 20.5 exon 21 p8 252 bp | TGAATGTGAGCCGAGTACAGGTAGGTCAGC<br>CAGAAGCTCTAATGCAGCTAGATCACTCCG<br>TGCCGCCCTTACATGGACATGGCGACTCAC<br>TTACACATTTACTCCTATCATCTTCATCTC<br>CTGTGTAGTTCACTCATAGATATGACCCTC<br>CCCTTCCTGCATCTTTCCTTCCCATTCTCC<br>CCCTTTCTTTAGCATTGTTAAAATTTATGT<br>GCTGTCATCCATCTCCAATCACTAG | 2 (19) | 0 (17) |
